# Validation of a quantitative PCR assay for the detection and quantification of the nematode Litylenchus crenatae in American beech leaf tissue

**DOI:** 10.64898/2026.09.03.749213

**Authors:** Eliana Torres-Bedoya, Andrew Miles, Justina Ker, Francesca Rotondo, Karen Snover-Clift, Pierluigi Bonello

## Abstract

Beech leaf disease (BLD), caused by the migratory plant pathogenic nematode, *Litylenchus crenatae* (LC, Anguinidae) is presently one of the most concerning forest diseases in the United States. BLD has spread rapidly across the landscape and is currently found in 15 U.S. states and Ontario, Canada, making monitoring a top priority. Here, we report on the development and testing of a probe-based qPCR detection assay that integrates previously reported LC-specific primers with a newly designed probe, enabling sensitive and reproducible quantification of nematodes in plant and non-plant matrices. This assay exhibited strong *in silico* and *in planta* specificity and high quantitative performance (R^2^ = 0.988–0.996), with amplification efficiencies of 90.0–96.8%. The assay also performed consistently on LC-spiked spore trap membranes used to monitor the fungal aerobiome, despite matrix-associated shifts in amplification efficiency. These results support the utility of the assay for monitoring and surveillance. The assay’s reliability was further evaluated by two independent diagnostic laboratories, demonstrating high inter-laboratory concordance in detection and strong agreement in relative quantification. The assay’s analytical sensitivity and reproducible performance align with key validation principles outlined by the APS Diagnostic Assay Validation Network (DAVN) and support its application as a molecular tool in experimental, diagnostic, and surveillance contexts, with important implications for national and international LC monitoring and disease management.

## Introduction

Beech leaf disease (BLD) is a rapidly expanding threat to forest ecosystems in North America, causing progressive canopy decline and widespread mortality in American beech (*Fagus grandifolia* Ehrh.) (Kantor et al., 2025; Shepherd et al., 2025). The disease is characterized by interveinal leaf banding, leaf thickening and crinkling, and bud abortion, with accompanying shifts in leaf microbiome, biochemistry, and physiology, which together reduce photosynthetic capacity, impair regeneration, and have long-term impacts on forest structure (Ewing et al., 2019; McIntire, 2023; Miles et al., 2026). Since its initial detection, BLD has spread extensively across the northeastern range of American beech, raising concerns about its ecological and economic costs.

The causal agent of BLD is the migratory nematode *Litylenchus crenatae* (LC, Anguinidae) (Carta et al., 2020; Vieira et al., 2023), an invasive foliar species (likely originating in Japan) that overwinters in dormant buds and colonizes developing leaves following budbreak (Wolf & Vieira, 2026). LC exhibits host specificity within the genus *Fagus*, and is pathogenic to *F. sylvatica* L. (European beech), *F. orientalis* Lipsky (Oriental beech), and *F. engleriana* Seeman ex Diels (Chinese or Engler’s beech), in addition to *F. grandifolia*, therefore representing a potential global threat (Carta et al., 2020; Kantor et al., 2025; Wolf & Vieira, 2024, 2026). Notably, disease severity varies widely among hosts, with the co-evolved *Fagus crenata* Blume (Japanese beech) exhibiting minimal damage when infected, suggesting it is resistant (Carta et al., 2020; Kanzaki et al., 2019), as expected for a co-evolved association (Wolf & Vieira, 2026). Additionally, LC is not the only nematode detected in beech tissues, other genera, including *Aphelenchoides* and *Plectus*, have also been reported (Reed et al., 2020).

Although American beech is not a major timber species, it plays a critical role in eastern North American forests, supporting wildlife and contributing to nutrient cycling and forest structure (Albright, 2018; McKenney-Easterling et al., 2000; Stephanson & Ribarik Coe, 2017). As a result, decline due to BLD poses significant risks to ecosystem stability and function. The impact of this disease extends beyond natural forests into urban landscapes and managed collections, where many ornamental beech species are susceptible. For example, arboreta such as The Morton Arboretum, Arnold Arboretum, the Holden Arboretum and Learning Gardens, and The Dawes Arboretum maintain valuable beech collections of horticultural, historical, and genetic significance, making BLD a major management concern in these settings.

Given the rapid spread and ecological importance of beech, effective monitoring and early detection are essential for disease management. Current evidence suggests that wind and rain contribute to nematode dispersal across the landscape (Fitza et al., 2024; Goraya et al., 2024). While tools such as spore impact traps that are deployed to capture fungal spores (Quesada et al., 2018) offer promise for monitoring environmental LC spread, accurate detection of LC still depends on reliable, reproducible, and precise diagnostic tools.

At present, LC detection relies on either microscopy or molecular approaches. Microscopy-based methods are labor-intensive, require expertise, and are unsuitable for high-throughput or quantitative applications. Molecular methods, including conventional PCR, quantitative PCR (qPCR), and recombinase polymerase amplification (RPA) (Burke et al., 2023; Kantor & Subbotin, 2026; Vieira et al., 2023), provide improved sensitivity and specificity but remain limited in important ways. For example, a qPCR assay targeting the ITS region (Burke et al., 2023) exhibited non-specific amplification and inconsistent performance in leaf tissue extracts under the conditions evaluated in this study; fatty acid- and retinol-binding gene (FAR)-based primers (Vieira et al., 2023), designed for purified nematode DNA, produce primer-dimer artifacts in complex plant matrices, compromising quantitative reliability; and conventional PCR and RPA-based methods (Kantor & Subbotin, 2026) do not provide quantitative estimates of nematode abundance.

Here, we present the development and validation of a FAR-based qPCR assay that incorporates a sequence-specific hydrolysis probe. This new assay enables quantification using purified nematode DNA, nematode-spiked, and naturally infested leaf and bud tissues, and experimentally spiked impact spore-trap membranes. We evaluated assay sensitivity, efficiency, linearity, matrix effects, and inter-laboratory reproducibility, and compared its performance with that of existing molecular protocols, following key principles outlined by the APS Diagnostic Assay Validation Network (DAVN) (Groth-Helms et al., 2023).

## Materials and Methods

### Plant material and nematode isolation

LC cannot currently be maintained or cultured *in vitro*, unlike certain other plant-parasitic nematode species. Therefore, for this study, wild reference nematodes were collected from both buds and leaves obtained in the Allegheny National Forest, Pennsylvania, within a plot network established and maintained as part of a long-term forest health monitoring program (Miles, 2023; Miles et al., 2026). During assay optimization, we used both fresh nematodes and nematodes preserved at -20° C, as they did not differ in DNA yield during processing. Naïve host tissue was used as a clean matrix for standardized spiking and extraction (i.e., for assay calibration and standard-curve development), collected in spring 2025 from American beech trees at The Ohio State University’s Chadwick Arboretum, a location outside the known zone of infestation. Naïve tissue was visually verified to be free of BLD symptoms and subsequently confirmed to be LC-negative using the probe-based qPCR assay.

Briefly, buds were dissected into individual scales and leaf primordia, then pooled into 15 mL conical tubes containing approximately 4 mL of molecular-grade water and 10 µL of Tween-20. In the case of leaves, ‘strips’ of the dark green interveinal tissue were extracted in 50 mL tubes in a similar manner. The tissue was vortexed for 5 min to dislodge nematodes, and the resulting suspension was filtered through a 25.4 µm mesh sieve. Extracts were centrifuged at 10,000 rpm for 5 min, after which the supernatant was discarded. Then 1 mL of water was added and the tube vortexed until the pellet was well-resuspended. The nematode suspension was then transferred to a 2.0 mL microcentrifuge tube. Depending on nematode density, samples were either re-centrifuged to concentrate the nematodes or diluted to facilitate nematode enumeration. Nematode counts were determined under a stereomicroscope; for counts exceeding 100 individuals, 10 µL volumetric subsampling was used to estimate total counts in triplicate, and the mean density of the suspension was calculated. Following extraction, LC suspensions were stored at -20 °C until further use.

### DNA extraction

DNA was extracted from leaf tissue using a modified two-day CTAB protocol (Clarke, 2009; Miles et al., 2026). Approximately 25 mg of frozen tissue was combined with five 2.4-mm metal beads in a 2.0 mL microcentrifuge tube, after which 980 µL of CTAB buffer (44 mM Tris, 56 mM Tris-HCl, 1.4 M NaCl, 50 mM Na_2_EDTA-H_2_O, 2.5% w/v CTAB, 1% w/v PVP-40) and 20 µL of 2-mercaptoethanol were added. Samples were vortexed for 1 min and incubated at 65 °C for 30 min with vigorous manual agitation every 10 min. Following incubation, 500 µL of chloroform:isoamyl alcohol (24:1) was added, the mixture was vortexed, and the samples were centrifuged at 14,000 rpm for 5 min. The aqueous phase was transferred to a new tube, mixed with an equal volume of cold isopropanol, and incubated at -20 °C overnight. The following day, samples were centrifuged at 14,000 rpm for 1 min, and the supernatant was discarded. Pellets were washed with 1 mL of 70% ethanol, air-dried, and resuspended in 50 µL of TE buffer and stored at -20 °C.

To build a standard curve, exact numbers of nematodes (0, 1, 10, 100, 1,000, 2,000) were added to 25 mg of naïve leaf tissue and extracted as described above. Both fresh and frozen nematodes were tested for spiking and yielded comparable extraction efficiencies and standard-curve performance. The same CTAB protocol was used for nematode-only samples. Procedures for spore traps are described further below.

### Primer validation and probe design

Given the novelty of this pathosystem and the absence of comprehensive public genomic or transcriptomic resources for LC, the FAR primers targeting the effector gene encoding a fatty acid- and retinol-binding protein (Vieira et al., 2023) were evaluated for potential off-target amplification. Primer sequences were queried against the NCBI nucleotide database using BLAST and against our in-house transcriptome assembly (Miles et al., in preparation) to evaluate predicted specificity.

To improve assay performance and eliminate secondary-structure and primer-dimer artifacts observed with SYBR Green chemistry (see below), a hydrolysis probe was designed within the FAR amplicon using Geneious Prime 2024.0.2 (https://www.geneious.com). The probe was included to enhance specificity and stability, and to support accurate quantification (**Figure S1**). Primer and probe sequences, as well as labeling chemistries, are listed in **Table 1**.

**Table 1.**
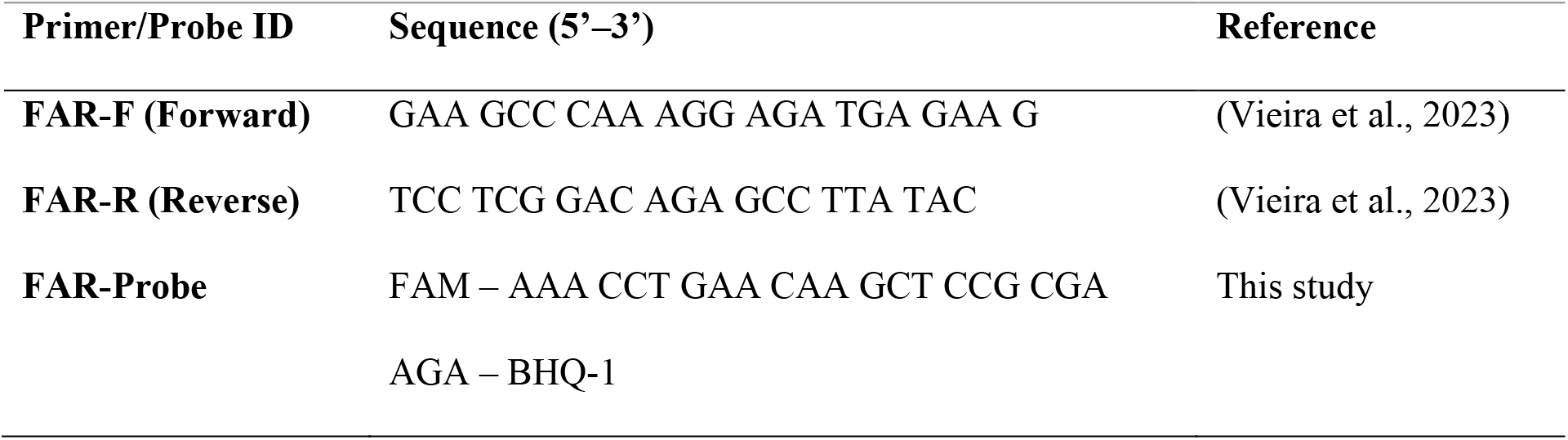
Sequences and labeling chemistries for the FAR-based qPCR assay.

### Assay specificity

Assay specificity was evaluated using a hierarchical approach: (1) determining whether naïve beech tissues contained any nematodes, (2) assessing primer and probe specificity to LC through *in silico* analysis, and (3) testing whether the assay amplified DNA from the commonly co-occurring non-target nematode *Aphelenchoides* spp.

To address the first question, a subset of naïve and LC-spiked leaf and bud samples was screened using conventional PCR targeting the nematode-specific D2-D3 expansion region of the 28S rRNA gene with primers D2A (5′-ACAAGTACCGTGAGGGAAAGTTG-3′) and D3B (5′-TCGGAAGGAACCAGCTACTA-3′) (Subbotin et al., 2006). Reactions were prepared with the GoTaq® G2 DNA Polymerase system (Promega, Madison, WI, USA) in a final volume of 25 µL. Negative controls included bacterial DNA, fungal DNA, and a no-template control containing molecular-grade water. Positive controls consisted of purified DNA from *Pratylenchus pseudocoffeae* (courtesy of the Lopez-Nicora Lab at Ohio State University; Consoli et al., 2026) and nematodes extracted from symptomatic beech leaves known to contain both LC and *Aphelenchoides* spp. (Reed et al., 2020). Thermal cycling conditions followed Subbotin et al. (2006). Amplicons were visualized on 1.8% agarose gels stained with SYBR Safe DNA Gel Stain (Invitrogen, Thermo Fisher Scientific, Waltham, MA, USA). Samples producing amplicons of the expected size (∼500-800 bp) were considered positive for nematodes.

The second question was addressed through an *in silico* specificity analysis of the primers and probe against representative annotated FAR sequences from plant-parasitic nematodes obtained from GenBank, including *Radopholus similis* (JN968974.1), *Meloidogyne incognita* (MF510388.1), *Aphelenchoides ritzemabosi* (KX816796.1), *Heterodera avenae* (KU877266.1), *Globodera pallida* (Y09293.2), and *Heterodera filipjevi* (KU877268.1). These sequences, together with LC, were aligned in SnapGene (GSL Biotech LLC, Chicago, IL, USA) using MUSCLE with 16 iterations, UPGMB clustering, and nucleotide sum-of-pairs scoring. Primers and the hydrolysis probe were mapped to the LC FAR sequence, and nucleotide mismatches between each non-target sequence and the corresponding primer- and probe-binding regions were quantified from the alignment.

To directly assess specificity against the naturally co-occurring non-target nematode *Aphelenchoides* spp. (third question), probe-based qPCR assays were conducted using DNA extracted from both LC and *Aphelenchoides* spp. The latter was provided by Christopher Taylor (The Ohio State University). Assays were performed using the optimized procedure described below to confirm the absence of non-target amplification from nematodes within the beech phytobiome.

### qPCR assay performance and validation

qPCR assay performance was evaluated in terms of analytical sensitivity, specificity, precision, and reproducibility following established diagnostic validation standards (Groth-Helms et al., 2023).

#### SYBR Green assay

qPCR amplification using the FAR primer pair was first evaluated with SYBR Green chemistry. Reactions were performed in a 10 µL volume containing 5 µL iTaq^TM^ Universal SYBR® Green Supermix (Bio-Rad Laboratories, Hercules, CA, USA), 0.3 µL each of forward and reverse primers (10 µM), 2.4 µL ultrapure water, and 2 µL of DNA template. Amplifications were carried out using a CFX Opus 96 Real-Time PCR System (Bio-Rad Laboratories). Thermal cycling consisted of an initial denaturation at 94 °C for 3 min, followed by 45 cycles of 95 °C for 10 s and 57 °C for 30 s. A melt-curve analysis from 65–95 °C in 0.05 °C increments was generated at the end of the run, and representative FAR amplicons were purified and submitted for Sanger sequencing to verify target identity through Ohio State’s Genomics Shared Resource. An annealing temperature gradient (55-65 °C) was evaluated using DNA from samples containing approximately 10 LC individuals, as well as water and LC leaf tissue controls.

#### Probe-based assay

qPCR reactions incorporating the hydrolysis probe were performed in a 10 µL volume containing 5 µL Luna® Universal Probe qPCR Master Mix (New England Biolabs, Ipswich, MA, USA), 0.3 µL each FAR primer (10 µM), 0.1 µL of the hydrolysis probe (10 µM), 2.3 µL of molecular-grade water, and 2 µL of DNA template. Thermal cycling conditions and instrumentation were identical to those used for the SYBR assay, with fluorescence acquisition at the end of each annealing step. Melt-curve analysis was not performed for probe-based reactions. An annealing temperature gradient (55-65 °C) was evaluated using DNA from samples containing approximately 10 LC individuals and LC-spiked leaf tissue (10 LC), as well as water no-template controls.

#### Quality control and calibration

For both SYBR Green and probe-based assays, water no-template controls and extraction blanks were included in every run. Two calibration approaches were used to evaluate assay performance: (1) DNA extracted from isolated nematodes (0, 1, 10, 100, 1,000, 2,000 individuals) and (2) naïve leaf tissue (25 mg) spiked with the same nematode quantities prior to extraction.

Each standard level was run in technical triplicate. Cq values were plotted against log_10_ (nematode quantity), and linear regressions were used to calculate the slope, efficiency, and R^2^ for each calibration method. DNA dilution effects were evaluated using undiluted, 1:10, 1:15, and 1:20 templates; an optimal 1:15 dilution was selected for all analyses based on amplification consistency and standard-curve performance.

Quantification accuracy was assessed by analyzing additional spiked samples containing nematode quantities not included in the standard curve and comparing measured values to actual counts.

#### Limit of detection (LOD)

The analytical limit of detection (LOD) was defined as the lowest nematode input that yielded successful amplification in ≥ 95% of reactions across independent extraction batches and qPCR runs, consistent with analytical sensitivity criteria commonly applied in diagnostic assay validation frameworks such as the APS DAVN. Detection was assessed using matrix-matched samples spiked with low nematode inputs (1 and 10 individuals), evaluated across four independent extractions, each analyzed in technical triplicate across 5 qPCR runs. Amplification success was scored based on the presence of a quantifiable Cq within the exponential phase of amplification.

#### Limit of quantification (LOQ)

The analytical limit of quantification was defined as the lowest nematode quantity that yielded reproducible quantification with a coefficient of variation (CV) < 25% across replicates and remained stable within the regression model, used here as the criterion for quantitative precision. LOQ determination was restricted to analytical conditions under controlled spiking and extraction procedures and does not account for additional variability associated with heterogeneous field-derived samples.

#### Inter-laboratory validation

Inter-laboratory reproducibility was assessed by implementing the probe-based qPCR assay in two independent plant disease diagnostic laboratories, Ohio State’s C. Wayne Ellett Plant and Pest Diagnostic Clinic and Cornell University’s Plant Disease Diagnostic Clinic, both of which are part of the National Plant Diagnostics Network, NPDN (npdn.org). Both laboratories received identical aliquots of DNA for standard curve construction; however, each laboratory independently prepared a 1:15 dilution of the supplied DNA for each standard concentration. Laboratories also received blind, spiked samples prepared and extracted in our laboratory. Participating laboratories used the same primer-probe concentrations, master mix, cycling conditions, and template input volumes.

Standard curve performance was compared between laboratories using analysis of covariance (ANCOVA) with the model Cq ∼ log_10_ (nematode quantity) × laboratory to evaluate differences in slope (interaction term) and intercept (main effect of laboratory).

For blinded samples, mean Cq values from technical triplicates were used to estimate log_10_-transformed nematode quantities based on laboratory-specific standard curves. Inter-laboratory agreement was assessed using Pearson correlation and Bland-Altman analyses performed on log_10_-transformed estimates, with calculation of mean bias and 95% limits of agreement. Qualitative concordance (detection vs no amplification) between laboratories was also recorded, allowing assessment of reproducibility in both detection and relative quantification, consistent with diagnostic validation approaches.

#### Bud tissue validation

To evaluate assay performance in a diagnostically relevant substrate, naïve whole buds were collected from American beech trees at the Chadwick Arboretum in Columbus, OH. Samples consisting of either one whole bud or a pool of five whole buds were spiked prior to extraction with defined numbers of LC (0, 1, 5, and 10 individuals). DNA was extracted using the same CTAB protocol described above for leaf tissue. Each treatment combination was evaluated across three independent qPCR runs with three technical replicates per run. Cq values and amplification success were recorded for each bud quantity and nematode input level.

#### Spore-trap validation

To evaluate assay performance in a different, inert matrix, spore-trap tapes (polyurethane membranes; Breathe-Easy^®^, Diversified Biotech, Dedham, MA, USA) (Quesada et al., 2018) were spiked with defined quantities of LC (1, 10, 100, 200, and 500 individuals). DNA was extracted using a modified CTAB protocol optimized for these membranes to minimize the effects of the polyurethane membrane’s chemical composition and physical structure, as well as its adhesive layer. Briefly, tapes were placed in 2 mL microcentrifuge tubes, flash-frozen in liquid nitrogen, and mechanically disrupted with a sterile microcentrifuge pestle. Subsequently, 800 µL of CTAB lysis buffer (100 mM Tris-HCl pH 8.0, 20 mM EDTA, 1.4 M NaCl, 2% w/v CTAB, 0.2% 2-mercaptoethanol) was added along with 25 mg PVP-40 (included at a higher proportion than in the leaf tissue protocol) to improve removal of potential PCR inhibitors associated with the polymeric and adhesive components of the matrix, and five sterile 2.4-mm metal beads. Samples were vortexed at maximum speed for 15 min and incubated at 65 °C for 25 min with periodic mixing.

Following incubation, 600 µL of chloroform:isoamyl alcohol (24:1) was added, samples were vortexed briefly, and centrifuged at 10,000 rpm for 15 min at 4 °C. The aqueous phase was transferred to a clean tube, and DNA was precipitated with 500 µL of cold isopropanol at -20 °C overnight. The following day, samples were centrifuged at 10,000 rpm for 15 min at 4 °C; the supernatant was discarded, and the pellets were washed with 70% ethanol. After complete evaporation of residual ethanol, DNA was resuspended in 40 µL molecular-grade water and stored at -20 °C until qPCR analysis.

Preliminary template-dilution tests in these samples indicated reduced amplification consistency at the lowest nematode input levels; therefore, DNA extracts from spore-trap samples were amplified without dilution. These samples were used exclusively to evaluate analytical performance and generate matrix-specific calibration curves in a non-plant substrate; field-collected spore-trap samples were not evaluated in this study.

### Comparison with other qPCR protocols

To contextualize the performance of the FAR-based probe assay developed in this study relative to existing molecular diagnostics, we compared it with the qPCR protocol employing primers 33F/234R targeting the ITS region (Burke et al., 2023). The assay was carried out as described by Burke et al. (2023), using the same DNA extracts evaluated with the FAR-based assay. Comparative assessments included amplification success across symptomatic and spiked leaf tissues, melt-curve profiles, presence of non-specific products, and the ability to generate standard curves under identical plate setups and technical replicate schemes.

## Results

### Primer validation and probe design

*In silico* analyses supported the predicted specificity of the FAR primer set for LC and the targeted FAR gene. No off-target hits were detected against our in-house *de novo* Trinity transcriptome or the NCBI RefSeq mRNA/RNA databases when restricted to Nematoda (taxid: 6231). Within our in-house LC transcriptome, two closely related putative FAR transcript isoforms were identified, both of which encompassed the primer and probe binding regions (**Table S1**). A single alignment was identified in the nucleotide collection for the entomopathogenic nematode *Oscheius tipulae*; however, this hit contained four mismatches in the forward primer and would produce a predicted amplicon of 3,871 bp, making amplification under qPCR conditions technically implausible.

Additional *in silico* analyses were restricted to Plants (taxid: 33090), *Fagus* (taxid: 21024), *Fagus grandifolia* (taxid: 60423), Fungi (taxid: 4751), and Bacteria (taxid: 2), and identified two partial alignments, primarily involving the forward primer, in *Fagus sylvatica* and *Quercus ilex*. These hits contained at least three mismatches and would generate predicted amplicons of approximately 340 bp in *Q. ilex* and ≥ 680 bp in *F. sylvatica*, well outside the size range compatible with efficient qPCR amplification.

The hydrolysis probe displayed thermodynamic characteristics compatible with efficient qPCR performance, with a GC content of 50% and a melting temperature of 68.3 °C. Predicted hairpin structures were weak (ΔG -0.2 to -1.1 kcal mol^-1^; Tm < 22 °C), indicating no intramolecular folding likely to interfere with hybridization. Homodimer formation was minimal, with the strongest predicted dimer involving only four paired bases (ΔG = -10.36 kcal mol^-1^). Because this interaction is short, internal, and not 3’-anchored, it is not expected to form under qPCR cycling conditions and remains within acceptable limits for hydrolysis probes. Heterodimer analysis likewise showed only weak, short complementarity between the probe and either primer, with all predicted interactions involving ≤ 4 contiguous bases, ΔG values < -6.24 kcal mol^-1^, and none at the primer 3’ end.

### Specificity assay

Conventional PCR targeting the D2–D3 expansion region of the 28S rRNA gene detected “generic” nematode DNA in a variety of samples (**Fig. S2**). Expected fragment size was observed with *P. pseudocoffeae* nematodes extracted from symptomatic beech tissue and all LC-spiked leaf and bud samples. Naïve leaf tissue also produced a clear amplicon, whereas naïve bud tissue did not produce any products, indicating presence of non-LC nematodes in leaf tissue but not in bud tissue. No amplification was observed with bacterial or fungal DNA or with the PCR blank.

The *in-silico* specificity analysis of six plant-parasitic nematode species with representative FAR homologs revealed that non-target species contained numerous nucleotide mismatches (**Fig. S3; Table S2).** The forward primer exhibited 5-11 mismatches, the hydrolysis probe 7-11 mismatches, and the reverse primer 5-13 mismatches relative to the homologous regions of the non-target FAR sequences. The greatest divergence was observed for both *Heterodera* species. The results indicate substantial sequence divergence between the LC-specific primers and the hydrolysis probe and homologous FAR sequences from non-target plant-parasitic nematode species.

As far as direct amplification of *Aphelenchoides* spp. is concerned, the probe-based assay consistently failed to amplify DNA of this nematode, and no amplification was observed in non-template controls. These results provide experimental support for the specificity of the probe-based assay against a non-target foliar nematode normally co-occurring in beech tissues.

### qPCR assay validation

#### SYBR Green assay

SYBR Green amplification using the FAR primers produced a detectable signal from LC DNA. However, while melt-curve analysis showed a distinct FAR-specific amplicon peak at 78.5–79 °C, it also consistently produced a lower-temperature peak at 71–73 °C, initially hypothesized to represent primer-derived amplification artifacts (**Fig. S4**). The lower-temperature peak appeared in nematode-only samples as well as in spiked and naturally infested leaf tissues and became prominent at low template concentrations. This recurrent artifact reduced sensitivity and compromised the quantitative reliability of the SYBR Green-based assay. To optimize amplification conditions and reduce melt-curve artifacts, a 55-65 °C annealing temperature gradient was performed using 1:15-diluted DNA samples containing approximately 10 LC individuals, along with water and American beech leaf DNA (**Fig. S5**). Both assays amplified the 10-LC leaf samples across much of the temperature gradient; however, increasing the annealing temperature delayed amplification and reduced endpoint fluorescence as a general trend. The SYBR assay also produced late amplification in water non-template controls across temperatures. Melt-curve analysis showed these reactions had a lower peak temperature, consistent with the formation of primer-derived non-specific products. The probe-based assay substantially reduced detectable non-specific fluorescence in water controls, consistent with the goal of increasing specificity. Naïve leaf DNA produced no consistent amplification in either assay, although an isolated late amplification was observed at the higher end of the gradient. Based on the temperature-gradient experiment, 57 °C was selected as the annealing temperature for the probe-based assay because it provided the best balance between efficient target amplification and suppression of non-specific signal. Under these conditions, the assay subsequently showed high amplification efficiency and linearity, consistent detection at low nematode loads, and no amplification of naïve leaf DNA, *Aphelenchoides* spp. DNA, or no-template controls.

Sanger sequencing of amplicons obtained from LC-containing samples confirmed amplification of the expected LC FAR target (**Table S3**). In contrast, sequencing of products associated with the lower-temperature melt peak in water controls yielded short sequences dominated by primer-derived regions, including sequence corresponding to the opposing primer, consistent with primer-dimer or closely related primer-derived amplification artifacts. These products lacked the internal sequence required for hydrolysis-probe binding, explaining why they contributed to SYBR Green fluorescence but were not detected by the probe-based assay.

#### Probe-based assay

The probe-based qPCR assay incorporating the FAR hydrolysis probe showed consistent and high-quality amplification across these runs using nematode-only DNA (**Fig. 1a**). Amplification curves showed clean baselines and uniform exponential phases with no detectable non-specific fluorescence, indicating suppression of the primer-derived non-specific signal observed in the SYBR Green assay (**Fig. S4)**.

**Figure 1.**
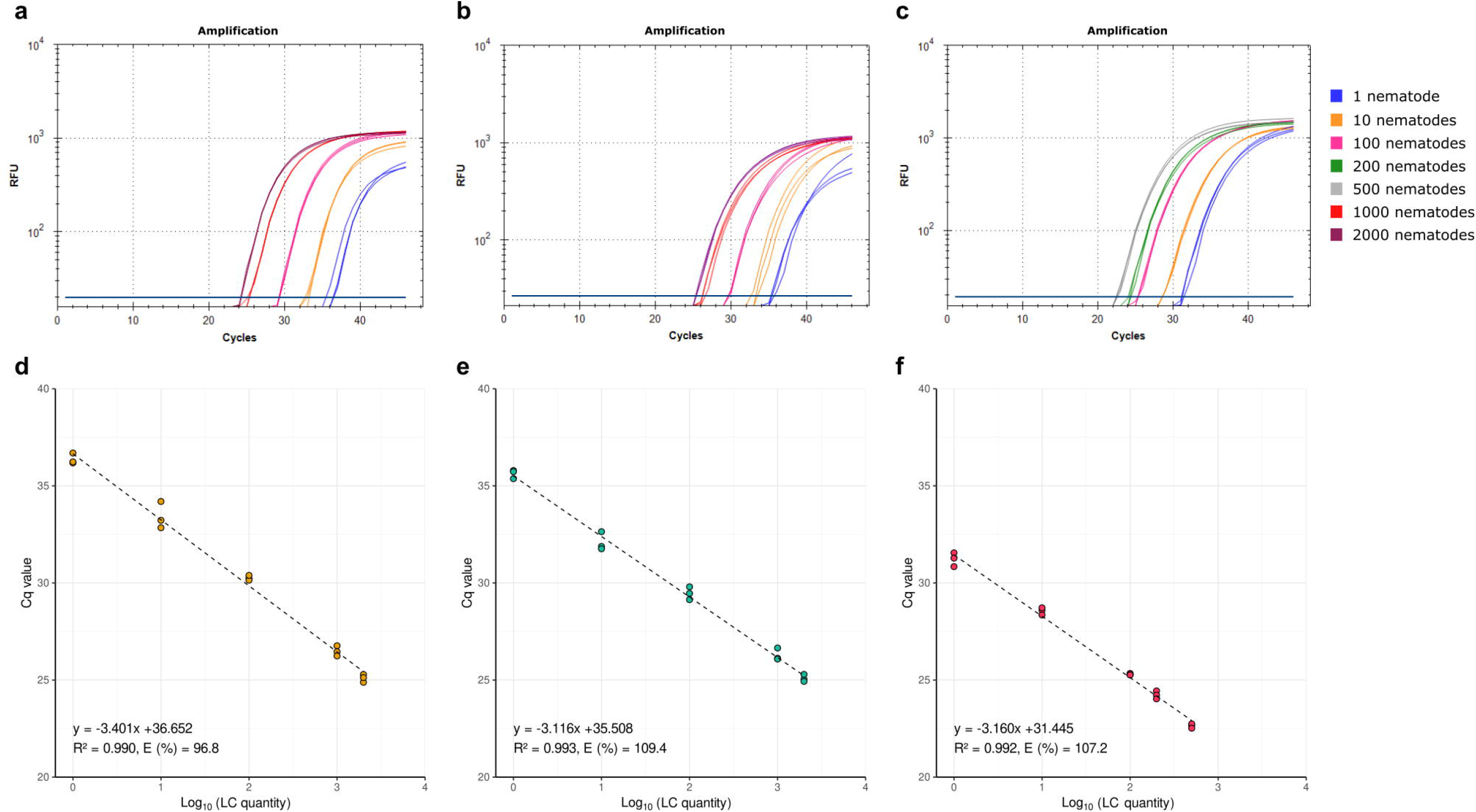
qPCR performance and standard curves for detection of LC across sample matrices. Amplification curves (a–c) and corresponding standard curves (d–f) were generated using (a, d) purified LC DNA, (b, e) naïve beech leaf tissue spiked with known numbers of LC prior to DNA extraction, and (c, f) spore-trap membranes spiked with known numbers of LC prior to DNA extraction. Amplification plots show fluorescence (RFU, log scale) across PCR cycles corresponding to increasing nematode quantities. Linear regressions were fit between Cq values and log_10_-transformed nematode quantities.

Standard curves generated from purified nematode DNA displayed strong analytical performance across four independent plates, with amplification efficiencies ranging from 90.0% to 96.8% and coefficients of determination (R^2^) between 0.990 and 0.994 (**Fig. S6**). A representative standard curve is shown in **Fig. 1d**. Cq values decreased predictably with each log_10_ increase in nematode quantity, and technical replicates exhibited low dispersion (CV 2.2–3.6%), indicating high repeatability (within-run precision) and stability across the tested dynamic range.

Residuals showed no evident change in dispersion across the fitted range, supporting the assumption of approximately constant variance across plates. Additionally, intercept variation was limited (34.78–36.65 Cq), consistent with stable amplification performance across plates.

Amplification was successful for all tested concentrations, including samples derived from a single nematode input, and Cq values at this level remained well within the range of exponential amplification. Replicate variability at the lowest input was low (CV = 2.2%), comparable to or lower than that observed at higher template quantities (CV = 2.6–3.6%). Accordingly, under the controlled conditions used here, amplification at the single-nematode level was consistently observed across independent runs and exhibited stable quantification precision, meeting the Minimum Information for Publication of Quantitative Real-Time PCR Experiments (MIQE) criteria (Bustin et al., 2009) for an analytical limit of detection of one nematode per reaction under nematode-only conditions, thereby demonstrating high analytical sensitivity under controlled conditions.

Although the analytical LOD is 1 nematode under the tested conditions, this value should be interpreted cautiously, as performance at low template abundance may be more susceptible to stochastic effects and matrix-associated variability in complex substrates such as leaf tissue. The LOQ, defined as the lowest concentration within the linear range of the standard curve with a coefficient of variation < 25%, was also 1 nematode per reaction under nematode-only conditions, reflecting analytical performance under controlled extraction and amplification conditions.

Standard curves generated from naïve leaf tissue spiked with known quantities of LC showed strong linear relationships between Cq and log_10_ (nematode quantity) across plates with R^2^ ranging from 0.988 to 0.996 (**Fig. 1b, 1e; Fig. S7**). Amplification efficiencies ranged from 105.9% to 110.7%, which fell within the range commonly observed for qPCR assays applied to complex biological matrices and are consistent with matrix-associated effects on reaction kinetics rather than non-specific amplification, particularly when template dilution is required to mitigate residual inhibitors. Importantly, efficiency estimates were reproducible across plates and showed stable slopes, high linearity, and low replicate variability (CV = 0.66–1.76%), indicating high precision and robustness across matrix-matched conditions.

The consistent amplification of single-nematode samples in both purified and spiked-tissue (mean Cq ≈ 35.3–35.6; CV = 0.64–1.57%) contexts establishes one nematode per reaction as the analytical LOD. Because Cq variance at this level was comparable to that at higher concentrations, the LOQ was also set to 1 nematode under the tested conditions, while recognizing that practical detection and quantification near this threshold may be influenced by sampling stochasticity, extraction efficiency, and matrix effects in field-derived samples.

#### Inter-laboratory reproducibility

Implementation of the probe-based FAR qPCR assay in two independent diagnostic laboratories, using the same PCR platform (CFX Opus 96, Bio-Rad Laboratories), yielded highly concordant qualitative results and strongly correlated quantitative estimates. Standard curves generated from identical DNA aliquots (independently diluted at each site) demonstrated comparable amplification efficiencies between laboratories (Lab A: 89.0%; Lab B: 81.1%) and strong linearity across the full dilution series (Lab A: R^2^ = 0.983; Lab B: R^2^ = 0.998; **Fig. 2a-b**). Analysis of covariance detected no significant difference in amplification efficiency between laboratories (log x lab interaction, p = 0.265) and no significant intercept shift (p = 0.154).

**Figure 2.**
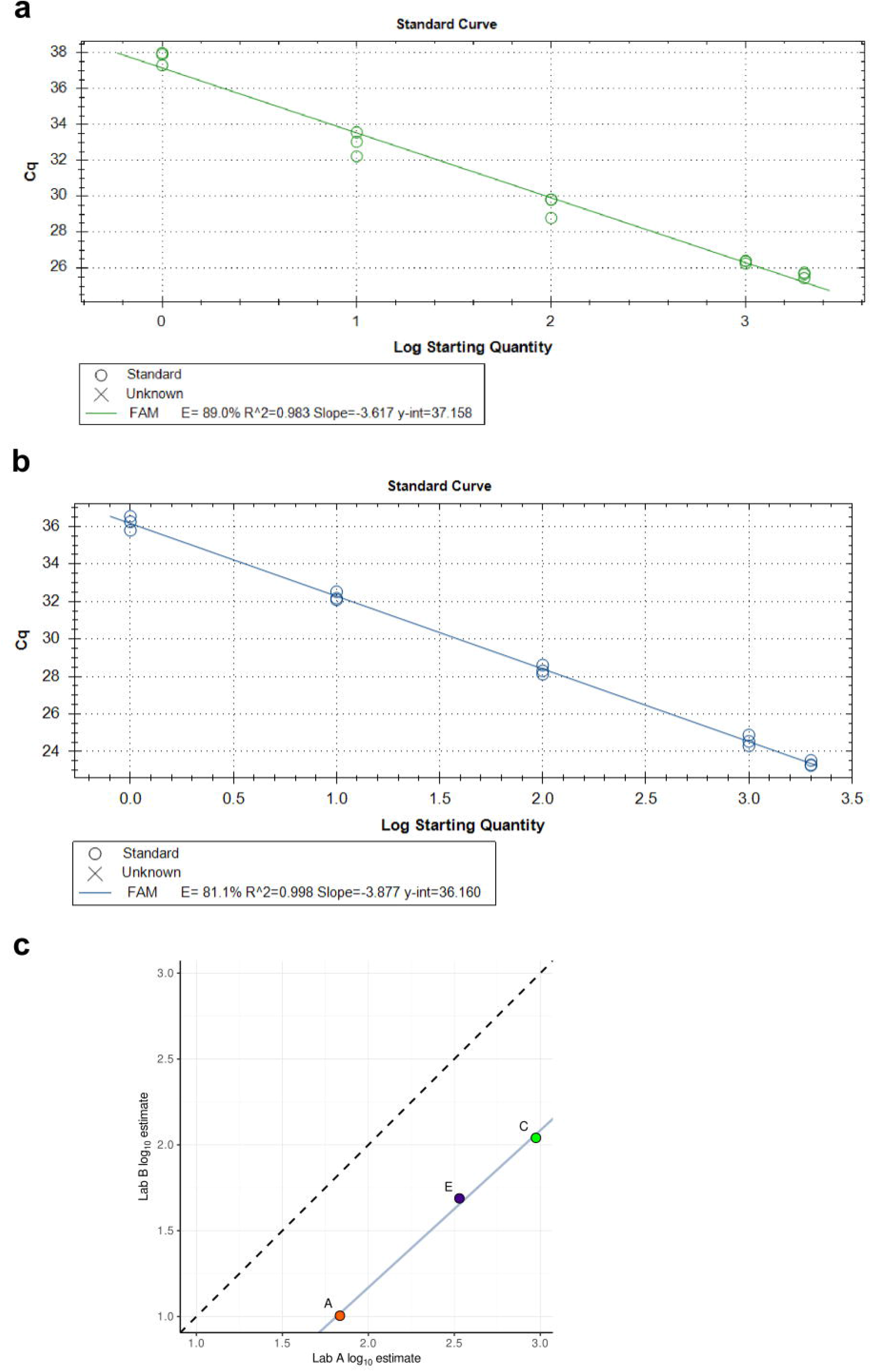
Inter-laboratory validation of the probe-based FAR qPCR assay. Standard curves generated independently by two diagnostic laboratories using identical DNA aliquots (with dilutions prepared at each site) are shown for (a) Laboratory A and (b) Laboratory B. Lines represent linear regression of Cq values against log_10_ starting quantity, with amplification efficiency (E), coefficient of determination (R^2^), slope, and intercept indicated. (c) Comparison of log_10_-estimated LC nematode quantities for positive blinded samples (a, c, and e) analyzed independently by both laboratories. The dashed line denotes the line of identity (y = x), and the solid line represents the fitted linear regression.

Blinded samples (A–E) were independently analyzed by both laboratories (**Table 2**). Qualitative detection was fully concordant: samples B and D were negative in both laboratories (no amplification observed), whereas samples A, C, and E were consistently detected at both sites. Among the positive blind samples, estimated log_10_ nematode quantities demonstrated near-perfect correlation (Pearson’s r = 0.998; Spearman’s ρ = 1.0; **Fig. 2c**). Bland-Altman revealed a consistent mean difference (bias) of 0.868 log units, with narrow 95% limits of agreement ranging from 0.757 to 0.980 log units. This corresponds to an approximately 7.4-fold systematic difference in absolute quantification between laboratories, indicating that although the assay was highly reproducible for detection and sample ranking, absolute quantitative estimates may vary between laboratories unless calibration procedures are further standardized. Low-abundance samples were consistently detected in both laboratories, and intra-laboratory Cq variability remained low across technical replicates (SD ≤ 0.54 cycles), supporting strong reproducibility in detection and relative quantification across laboratories.

**Table 2.** Inter-laboratory comparison of blinded samples (A–E) analyzed using the probe-based FAR qPCR assay. Mean Cq values (± SD, technical triplicates) are shown for each laboratory. Log_10_ quantities were calculated using laboratory-specific standard curves. Δ log_10_ represents the difference between laboratories (Lab A–Lab B), while fold diff corresponds to 10^Δ log_10_. Samples B and D showed no amplification (NA) in either laboratory.

| Sample | Cq Lab A | Cq Lab B | Log10 Lab A | Log10 Lab B | $\Delta$ log10 | Fold diff |
| --- | --- | --- | --- | --- | --- | --- |
| **A** | 29.5 ± 0.41 | 32.3 ± 0.11 | 1.83 | 1.00 | 0.83 | 6.76 |
| **B** | NA | NA | NA | NA | NA | NA |
| **C** | 25.3 ± 0.21 | 28.2 ± 0.10 | 2.97 | 2.04 | 0.93 | 8.59 |
| **D** | NA | NA | NA | NA | NA | NA |
| **E** | 26.9 ± 0.49 | 29.6 ± 0.54 | 2.53 | 1.69 | 0.84 | 6.94 |

#### Bud tissue validation

Detection of LC was evaluated in both single-bud and pooled-bud matrices containing defined nematode loads. No amplification was observed in negative controls (0 nematodes) for either matrix (**Table S4**). In single-bud samples, amplification was consistently detected at inputs of 1, 5, and 10 nematodes across replicate reactions, with mean Cq values generally decreasing as nematode input increased. In contrast, detection in the pooled five-bud matrix was less consistent at the lowest input level. While samples containing five or ten nematodes in five bud-pools produced consistent amplification across replicates, samples containing a single nematode in five buds showed sporadic amplification (2/9 reactions) or failed to amplify (**Table S4**). These observations are consistent with a practical detection threshold of approximately one nematode per bud under the extraction and assay conditions used in this study.

#### Spore-trap validation

Matrix-specific calibration using spore-trap tapes spiked with known quantities of LC demonstrated consistent quantitative performance of the probe-based FAR qPCR assay in this non-plant substrate. Standard curves generated from four independent plates showed strong linear relationships between Cq and log_10_ (nematode quantity), with R^2^ values ranging from 0.988 to 0.992 and amplification efficiencies between 107.0% and 109.0% (**Fig. 1c,f; Fig. S8**). As observed for plant-derived extracts, efficiencies slightly above 100% are consistent with matrix-associated effects in non-biological substrates. Slopes were highly consistent across plates (-3.13 to -3.17), indicating stable amplification kinetics despite the distinct chemical and physical properties of the adhesive tape matrix.

Technical replicate variability was low across all concentration levels, with coefficients of variation ranging from 0.56% to 1.98% within plates and 0.69% to 1.41% when pooled across plates, indicating high repeatability and robustness in this matrix. Mean Cq values decreased predictably with increasing nematode input, and reactions containing the lowest spiked levels were consistently amplified on every plate (mean Cq ≈ 31.2–31.4) when undiluted DNA samples were used, demonstrating reliable detection near the lower quantitative range in this environmental matrix. No increase in dispersion or deviation from linearity was observed relative to purified nematode or plant-derived extracts, indicating that the spore-trap substrate did not introduce measurable inhibition or stochastic amplification effects.

### Comparison with a previously published qPCR protocol

Performance of the probe-based FAR qPCR assay was compared with a previously published ITS-based qPCR protocol using primers 33F/234R when applied to naïve leaf tissue spiked with defined quantities of LC. Under the original reaction conditions described for the ITS assay, amplification was inconsistent across template concentrations, and melt-curve analysis revealed complex profiles with multiple peaks and broad peak shapes, indicating substantial non-specific amplification in plant-derived extracts **(Fig. 3**). These artifacts were evident in both standard dilutions and naturally infested tissue samples, thereby compromising quantitative interpretability under the tested experimental conditions.

**Figure 3.**
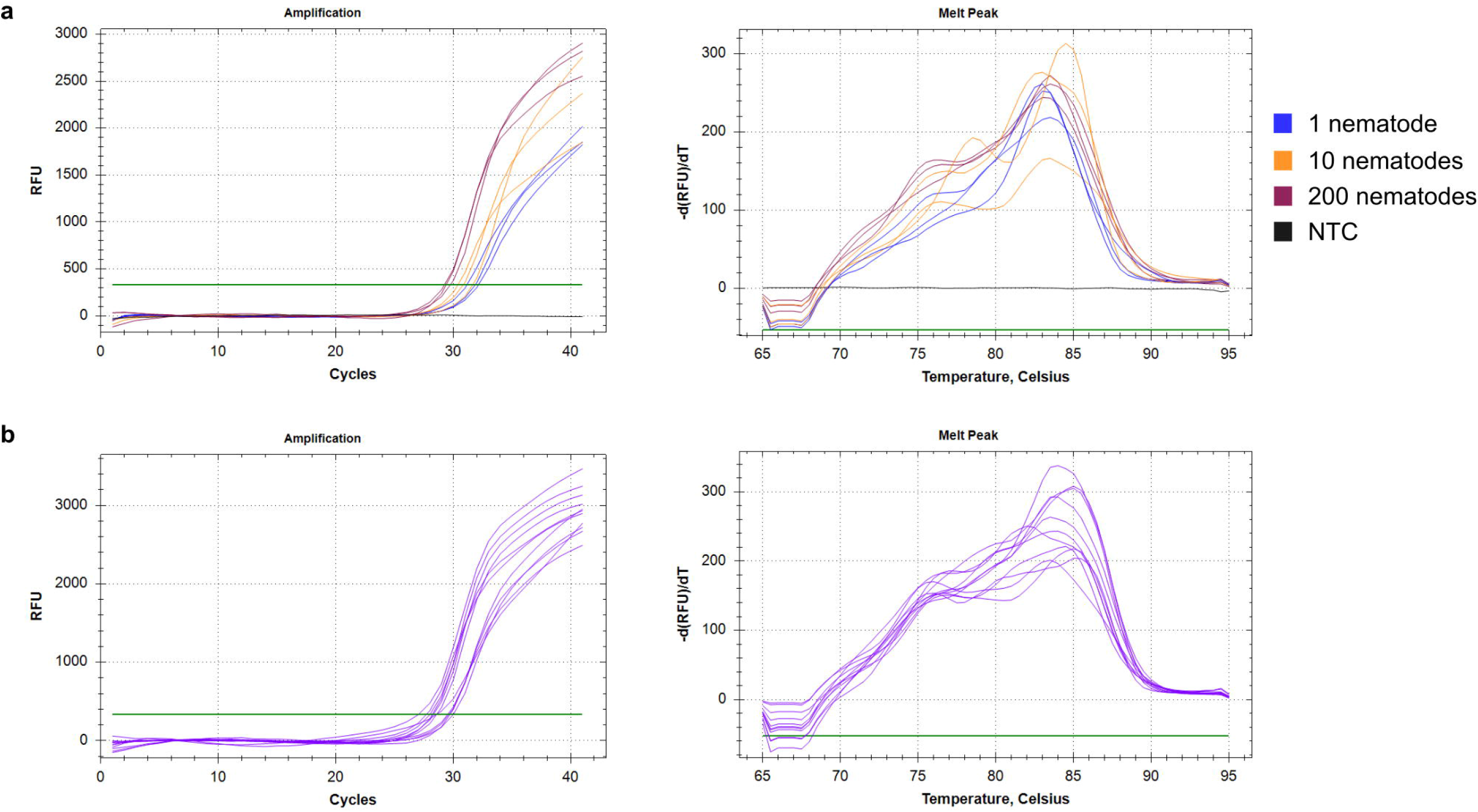
Performance of the ITS-based qPCR assay. Amplification curves and melt-curve profiles obtained from (a) naïve beech leaf tissue spiked with known quantities of LC and (b) naturally infested leaf samples.

To assess whether assay performance could be improved through protocol modification, two adjustments were evaluated: omitting the extension step (72 °C for 60 s) and increasing the annealing temperature by 1 °C (from 58 °C to 59 °C). Removal of the extension step resulted in minor changes to amplification kinetics but did not resolve the presence of multiple melt peaks or reduce non-specific signal accumulation (**Fig. S9**). Increasing the annealing temperature improved overall amplification stringency; however, melt-curve profiles continued to exhibit overlapping peaks and incomplete separation between the expected amplicon and non-specific products (**Fig. S10**).

Across tested conditions, the ITS-based assay did not yield reproducible standard curves with a single, well-defined melt peak when applied to plant tissue extracts, thereby limiting its suitability for quantitative applications in plant matrices. In contrast, the probe-based FAR assay produced clean amplification profiles and consistent quantitative performance across spiked tissue samples and naturally infested leaf material (**Fig. 4**), supporting an advantage of probe-mediated detection for suppressing non-specific signal in complex plant matrices under the conditions evaluated in this study.

**Figure 4.**
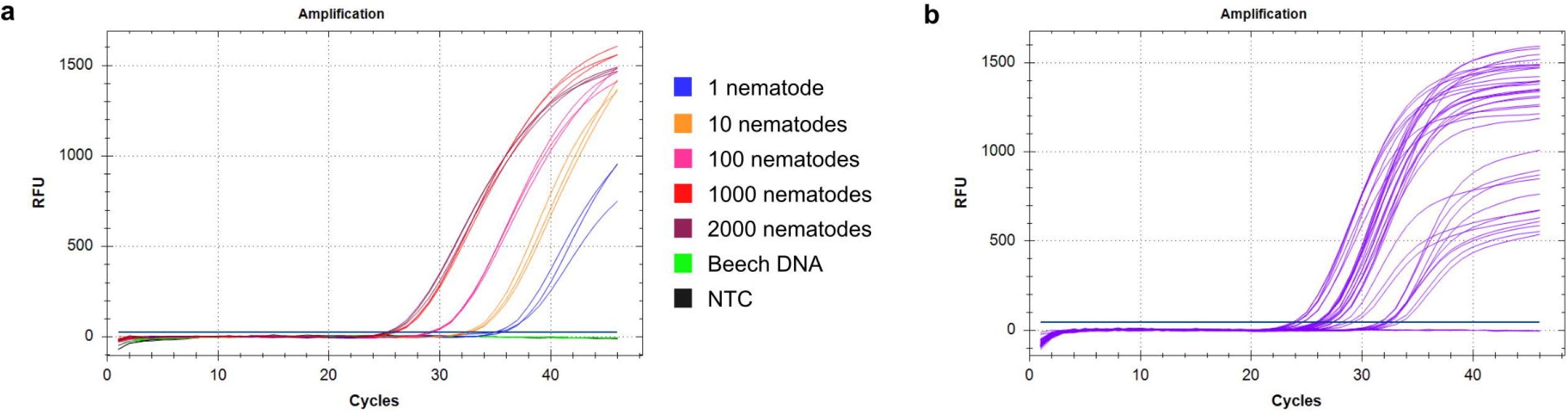
Performance of the probe-based FAR qPCR assay developed in this study. Amplification curves obtained from (a) naïve beech leaf tissue spiked with known quantities of LC and (b) naturally infested leaf samples.

## Discussion

In this study, we designed and validated a probe-based qPCR assay targeting a nematode-specific FAR gene to enable sensitive, selective, and reproducible quantification of LC from purified nematode DNA, plant-derived extracts, and spore-trap tape samples. By integrating a hydrolysis probe into a previously reported approach (Vieira et al., 2023), this assay suppressed non-specific signal and primer-dimer artifacts that limit the usefulness of other existing approaches, thereby improving quantitative interpretability in complex substrates, particularly at low nematode abundance and in plant-derived extracts. Annealing-temperature optimization further demonstrated that 57 °C provided an appropriate balance between target amplification and specificity for the primer-probe system, with assay performance subsequently supported by independent laboratory validation. This is critical, as reliable quantification of LC is a prerequisite for advancing research on BLD, including infection dynamics, spatio-temporal variation in nematode abundance, and mechanisms of dispersal.

A persistent challenge in molecular diagnostics across diverse pathosystems is selecting molecular targets that balance biological relevance with taxonomic resolution. FAR proteins (FARs) represent a relevant class of nematode-associated proteins with high affinity for fatty acids, retinol, and retinoic acids (Kennedy et al., 2013; Vieira et al., 2017). Their biological relevance stems from the inability of nematodes to synthesize fatty acids *de novo*, requiring lipid acquisition from the host or the surrounding environment (Kennedy et al., 2013; Vieira et al., 2017). FAR proteins have been implicated in lipid binding and sequestration during host-parasite interactions, supporting their involvement during infection (Kennedy et al., 2013; Vieira et al., 2017). Although FAR homologs are not exclusive to nematodes, the evolutionary history of this gene family remains incompletely resolved (Yuan et al., 2021). The FAR primer pair designed by Vieira et al. (2023) demonstrated strong predicted specificity for LC and provided sufficient discriminatory capacity for quantitative detection under the experimental conditions validated here.

Analytical validation using raw nematode DNA demonstrated strong linearity, high amplification efficiency, and low technical variability across independent plates, consistent with MIQE guidelines for qPCR assays (Bustin et al., 2009). The assay consistently amplified reactions containing a single nematode, with variance comparable to that observed at higher template inputs, supporting high analytical sensitivity and repeatability under controlled conditions. However, the complex biological context of the phytobiome, including variation in nematode size (resulting in variable FAR copy numbers/nematode), may affect quantification accuracy. Additionally, this study did not evaluate the detectability or quantitative contribution of nematode eggs, which are found throughout host tissues (Carta et al., 2020; Loyd et al., 2024; Vieira et al., 2023). Future refinements may also benefit from approaches that distinguish DNA from viable versus non-viable nematodes. For example, propidium monoazide treatment prior to extraction has been used in other systems to suppress amplification of DNA from dead cells and could be leveraged in situations where quantification of viable specimens is desirable (Christoforou et al., 2014; Sert Çelik et al., 2020).

Although the performance of our assays meets formal criteria for analytical LOD and LOQ of 1 nematode per reaction under nematode-only conditions, these thresholds should be interpreted cautiously when applied to complex matrices, such as the buds, leaves, and spore tapes used in this study. At very low template concentration, stochastic sampling effects, variability in extraction efficiency, and matrix-specific inhibition can disproportionately influence amplification behavior (Taylor et al., 2019), emphasizing the need for matrix-specific validation. Accurate standard-curve construction is therefore essential for maintaining quantitative reliability, particularly for microscopic organisms such as LC, where visual detection, enumeration, and transfer of individual nematodes are inherently challenging and can introduce substantial variability if not rigorously controlled.

Standard curves generated from naïve leaf tissue spiked with known quantities of LC retained strong linearity and low variability, despite a modest upward shift in apparent amplification efficiency. Such shifts are well documented in qPCR assays applied to complex biological matrices and likely reflect subtle alterations in reaction kinetics arising from residual inhibitors, changes in template accessibility, or differences in effective reaction volume following dilution (Bustin et al., 2009; De Chaves et al., 2023; Svec et al., 2015; Taylor et al., 2019). Nevertheless, slopes and intercepts remained highly consistent across plates, and replicate variability remained low even at the single-nematode level, indicating stable quantitative behavior across the assay’s dynamic range. Collectively, these results demonstrate that this FAR probe-based assay is resistant to plant matrix effects and supports reliable quantitative applications in foliar tissues when calibration is performed using matrix-matched standards.

Inter-laboratory validation further supports the practical utility of this assay in diagnostic settings. Standard curves generated independently by two University-supported plant disease diagnostic laboratories showed comparable amplification efficiencies, strong linearity, and ultimately overall concordance of LC quantification in blind samples between the two laboratories. However, the Bland-Altman analysis revealed a systematic difference in absolute quantification between laboratories. Thus, while the assay appears robust for qualitative detection and relative comparisons among samples within a laboratory, absolute nematode estimates may remain laboratory-dependent unless standard preparation, dilution handling, and calibration procedures are further harmonized, highlighting the distinction between reproducibility of detection and relative quantification versus absolute measurement agreement.

Although qualitative concordance between laboratories was high, formal estimates of diagnostic sensitivity and specificity were not calculated, as this study was based on controlled spiking experiments rather than field-derived samples with independently verified infection status.

In addition to quantitative performance, specificity in the presence of non-target nematode DNA is critical for reliable application in plant-derived matrices. Given that LC has not been successfully cultured *in vitro*, we conducted an indirect specificity assay alongside an *in-silico* alignment, which provided supporting evidence for the selective detection of LC by our assay. Conventional PCR targeting the D2–D3 region confirmed the presence of nematode DNA in both LC-spiked samples and naïve leaf tissue, demonstrating that plant-derived extracts can contain background nematode communities independent of experimental inoculation. In contrast, naïve bud tissue did not yield detectable amplification, suggesting lower or absent background nematode presence in this tissue under the conditions tested.

Importantly, despite the detection of non-target nematode DNA in naïve leaf tissue, no amplification was observed with our assay in these samples, indicating that the assay does not cross-react with non-LC nematodes present in the phytobiome. Furthermore, alignment with other plant-parasitic nematodes revealed significant mismatches across the primers and probe. These findings provide matrix-based support for assay selectivity in biologically relevant substrates and reinforce its suitability for applications in which complex, potentially confounding DNA backgrounds are expected, such as field-derived leaf samples and environmental surveillance. However, broader empirical testing against a wider range of non-target nematodes would further strengthen confidence in analytical specificity across diverse diagnostic contexts.

Taken together, the validation framework applied here aligns with key principles outlined by the APS DAVN (Groth-Helms et al., 2023), including the evaluation of analytical sensitivity, specificity, precision, and inter-laboratory reproducibility under controlled and matrix-relevant conditions. It also provides a valuable example of how research programs can work directly with the NPDN. Diagnosticians rely on research programs to develop and validate procedures, ensuring accurate test methods for identifying various target organisms. Using the NPDN diagnostic laboratories at Ohio State University and Cornell University helped demonstrate that this assay could be implemented successfully in diagnostic settings, benefiting both research and extension services and further supporting DAVN validation principles.

The analyses performed here address components of DAVN Tier 1 analytical validation and includes interlaboratory evaluation consistent with Tier 3 validation principles, but more extensive exclusivity testing would be required for higher-tier validation and broader diagnostic deployment. Given the currently reported low genetic variability of LC, inclusivity testing may be of lower priority, as evidenced by 100% sequence similarity at the LSU and ITS loci across populations (Fitza et al., 2024), a pattern consistent with a founder effect following a single introduction event in the invaded region. Herein, we expanded exclusivity testing to further support analytical specificity by including non-target nematodes, including foliar *Aphelenchoides* spp. isolated directly from American beech tissues. Future experimental testing could include additional plant-parasitic nematodes and environmentally ubiquitous taxa.

An additional validation step evaluated assay performance in bud tissue, which represents the substrate most submitted for diagnostic evaluation of BLD. Under the controlled spiking conditions used here, the assay consistently detected LC when a single nematode was present in an individual bud, demonstrating that the assay retains high sensitivity in this biologically relevant matrix. When nematodes were distributed within pooled bud tissue, however, amplification at the lowest input level became less consistent, whereas samples containing five or more nematodes in pooled buds amplified reliably. This pattern likely reflects the combined influence of dilution effects and stochastic sampling when very low nematode numbers are distributed across larger tissue volumes, rather than a limitation of the assay itself. These observations highlight the importance of considering the tissue matrix and sampling scale when interpreting detection thresholds in practical diagnostic contexts, and they support the use of bud tissue as a suitable substrate for molecular detection of LC. For quantitative applications involving bud tissue, matrix-matched calibration curves generated from bud extracts may be considered, as extraction efficiency and the presence of inhibitory compounds can vary among plant tissues.

Additionally, comparison of individual- and pooled-bud samples provides practical guidance for selecting an appropriate processing strategy for diagnostics. Although pooling can increase sample processing efficiency in clinical or experimental settings, detection was less consistent at the lowest infestation level, equivalent to one nematode per bud, indicating that negative results from pooled samples should be interpreted cautiously. In contrast, LC was consistently detected in individual buds containing a single nematode, demonstrating that the assay retains high sensitivity within a diagnostically relevant plant matrix, under certain processing conditions.

Beyond host tissues, the successful amplification and quantification of LC from experimentally spiked spore-trap membranes supports the potential application of this assay to non-plant substrates and provides a basis for future evaluation of spore traps as environmental surveillance tools. Of particular note, spore-trap tapes are a challenging matrix because adhesive compounds can inhibit PCR amplification or increase stochastic variability. Despite these constraints, the FAR assay yielded highly consistent standard curves across independent plates, with strong linearity, stable slopes, and low coefficients of variation across all tested concentrations. The absence of increased dispersion or deviation from linearity relative to plant-derived or raw nematode samples indicates that the spore-trap substrate did not cause measurable inhibition or amplification instability under the selected extraction and amplification conditions.

Comparison with a previously published ITS-based qPCR protocol highlights the practical advantages of probe-mediated detection in complex plant matrices. The ITS assay performed inconsistently when applied to both spiked and naturally infested beech tissue, with complex melt-curve profiles indicative of non-specific products. Protocol modifications modestly increased stringency but failed to eliminate these artifacts or restore reliable quantitative performance.

In addition to laboratory-based quantitative assays such as the one developed here, recent work has shown that RPA combined with lateral flow dipsticks (LF-RPA) can support rapid field detection of LC (Kantor & Subbotin, 2026), highlighting the growing need for complementary diagnostic tools adapted to different surveillance and management contexts and needs.

In conclusion, the probe-based FAR qPCR assay described here offers a sensitive, selective, and versatile eDNA platform for quantifying LC across diverse biological and environmental matrices. Its robust analytical performance, coupled with demonstrated inter-laboratory consistency in detection and sample ranking, positions this assay as a foundational tool for advancing research, diagnostics, and surveillance of beech leaf disease.

## Acknowledgements

This project was funded by USDA Forest Service Joint Venture Agreement 24-JV-11242314-012 and USDA Forest Service Cooperative Agreement 23-DG-11094200-485 with Ohio State University. Additional funding included grants to AM from the Kurt Gottschalk Science Fund (Society of American Foresters) and the Infectious Diseases Institute, Host Defense and Microbial Biology New Techniques Grant Program (Ohio State).

## Data availability statement

Additional data that supports the findings of this study are available from the corresponding author upon request.

