## Supplemental for "Validation of a quantitative PCR assay for the detection and quantification of the nematode Litylenchus crenatae in American beech leaf tissue"

**Supplementary Tables**

**Table S1.** Nucleotide sequences of the two LC FAR transcript isoforms identified in the in-house transcriptome assembly (unpublished). The FAR-F forward primer is highlighted in yellow, the FAR-R reverse primer in magenta, and the hydrolysis probe in green.

| **Transcript ID** | **Sequence** |
| --- | --- |
| TRINITY_DN5016_c0_g2_i4.p1 | ATGACTGAGGAAGACAAGACCGTGCTCAAGGACTTGGCCGGCCAGCATGCTAGCTTTGAGAACGAGGAGCAGGCTTTGAATGCCCTTAAAGAGAAGAGCCCAAAACTTTACGAGAAGGCTAAGGCTCTGCACGAGCTGGTCAAGAACAAGATCAACGAGCTCACAAACGCTGATGCCAAGACCTTTGTCAACAACATTATCGCCAAACTGCGTGCCTTGAAGCCCAAAGGAGATGAGAAGCCAAACCTGAACAAGCTCCGCGAAGAGGCCAACAACATCATCAATGAGTATAAGGCTCTGTCCGAGGAAGCCAAGGAGAACCTGAAGGCCACTTTCCCCAAAATTACCGGAGTAATCCAGAATGAGAAGTTCCAAAAATTGGCTAAAGGCTTGCTCAAGACTGACGCTCCTGCTGCCTAA |
| TRINITY_DN5016_c0_g2_i7.p1 | ATGTTGATGCGCTGCGTCGTGTTGCTTGCCCTGATGATTGTATACGTTAATGCCAATGCCCTCCCCACCTTTAACTTTAACCAGATCCCGGAGCAGTTCAAAGATATTGTACCTGAAGAAGTGAAAAAATTCTATGATGAGATGACTGAGGAAGACAAGACCGTGCTCAAGGACTTGGCCGGCCAGCATGCTAGCTTTGAGAACGAGGAGCAGGCTTTGAATGCCCTTAAAGAGAAGAGCCCAAAACTTTACGAGAAGGCTAAGGCTCTGCACGAGCTGGTCAAGAACAAGATCAACGAGCTCACAAACGCTGATGCCAAGACCTTTGTCAACAACATTATCGCCAAACTGCGTGCCTTGAAGCCCAAAGGAGATGAGAAGCCAAACCTGAACAAGCTCCGCGAAGAGGCCAACAACATCATCAATGAGTATAAGGCTCTGTCCGAGGAAGCCAAGGAGAACCTGAAGGCCACTTTCCCCAAAATTACCGGAGTAATCCAGAATGAGAAGTTCCAAAAATTGGCTAAAGGCTTGCTCAAGACTGACGCTCCTGCTGCCTAA |

**Table S2.** Number of nucleotide mismatches between the LC-specific qPCR assay primers and probe and homologous regions of representative plant-parasitic nematode FAR sequences.

| Species/Target | Forward Primer (22 Bp) | Far Probe (24 Bp) | Reverse Primer (21 Bp) |
| --- | --- | --- | --- |
| LC FAR-1 target | 0 | 0 | 0 |
| *Aphelenchoides ritzemabosi* | 9 | 10 | 5 |
| *Globodera pallida* | 8 | 9 | 12 |
| *Heterodera avenae* | 11 | 11 | 13 |
| *Heterodera filipjevi* | 11 | 11 | 12 |
| *Meloidogyne incognita* | 8 | 7 | 12 |
| *Radopholus similis* | 5 | 9 | 13 |

**Table S3.** Sanger sequences of representative FAR amplicons generated from water controls, LC-containing SYBR Green reactions, and LC-containing probe-based qPCR reactions. Trimmed high-quality reads were compared with the expected *Litylenchus crenatae* FAR amplicon to evaluate target amplification and characterize primer-derived non-specific products.

| **Sample** | **Read** | **Trimmed sequence (5′–3′)** | **Interpretation** |
| --- | --- | --- | --- |
| Water control product | Forward | CATCATCAATGAGTATAAG  GCTCTGTCCGAGGA | Primer-derived sequence; terminal region corresponds to reverse-primer sequence |
| Water control product | Reverse | GTTCAGGTTTGGCTTCTCA  TCTCCTTTGGGCTTCAATG | Primer-derived sequence; contains sequence corresponding to the forward-primer region |
| SYBR product | Forward | CATCATCAATGAGTATAAG  GCTCTGTCCGAGG | Matches the expected LC FAR amplicon across the full trimmed read - Target amplification confirmed |
| SYBR product | Reverse, reverse-complemented | AAGCCCAAAGGAGATGA  GAAGCCAAACCTGAACAA | Matches the expected LC FAR amplicon across the full trimmed read - Target amplification confirmed |
| Probe-based product | Forward | CATCATCAATGAGTATAA  GGCTCTGTCCGAGGAA | Matches the expected LC FAR amplicon across the full trimmed read - Target amplification confirmed |
| Probe-based product | Reverse, reverse-complemented | AAGCCCAAAGGAGATG | Short high-quality read matching the expected LC FAR amplicon across the full trimmed region - Supports target amplification |

**Table S4.** Detection of LC in spiked beech bud tissue across independent qPCR runs. Cq values obtained from three independent qPCR plates with three technical replicates per plate are shown for samples containing defined numbers of nematodes in either a single bud or a pool of five buds. Numbers in parentheses indicate the number of nematodes added to each sample prior to DNA extraction. “-” indicates no amplification was detected.

| Plate | Replicate | 1 Bud (0) | 1 Bud (1) | 1 Bud (5) | 1 Bud (10) | 5 Buds (0) | 5 Buds (1) | 5 Buds (5) | 5 Buds (10) |
| --- | --- | --- | --- | --- | --- | --- | --- | --- | --- |
| 1 | 1 | - | 32.5 | 31.9 | 30.3 | - | - | 35.4 | 31.9 |
|  | 2 | - | 33.0 | 32.7 | 30.1 | - | 35.1 | 34.2 | 31.3 |
|  | 3 | - | 32.0 | 31.6 | 30.0 | - | - | 34.0 | 32.1 |
| 2 | 1 | - | 35.1 | 33.8 | 30.8 | - | - | 33.0 | 34.3 |
|  | 2 | - | 35.2 | 33.6 | 31.5 | - | - | 36.1 | 33.2 |
|  | 3 | - | 35.8 | 33.2 | 31.3 | - | - | 38.1 | 34.8 |
| 3 | 1 | - | 33.1 | 34.9 | 31.2 | - | - | 36.9 | 33.2 |
|  | 2 | - | 33.4 | 34.4 | 31.2 | - | - | 37.3 | 33.8 |
|  | 3 | - | 34.1 | 34.5 | 31.7 | - | 44.9 | 35.1 | 33.3 |

**Supplementary Figures**

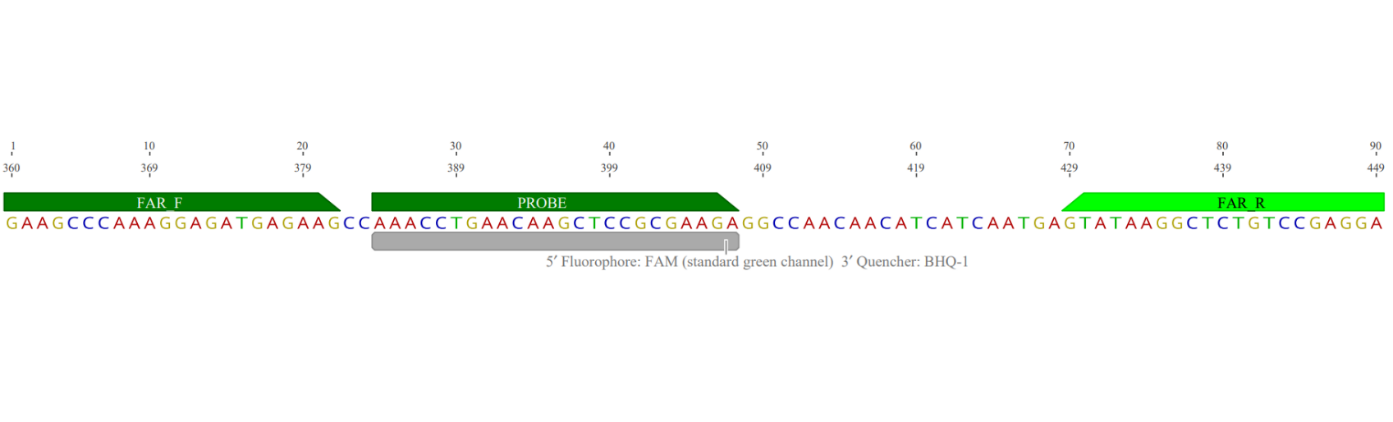
**Figure S1.** FAR amplicon structure and primer–probe alignment. Schematic representation of the FAR amplicon targeted by the qPCR assay, showing the relative positions and orientations of the FAR-F forward primer, FAR-R reverse primer, and the internal hydrolysis probe.

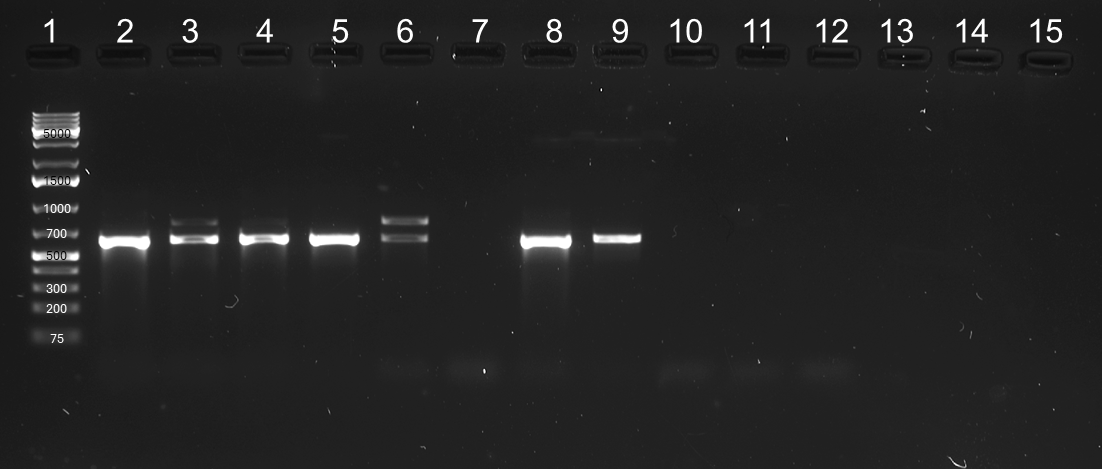

**Figure S2.** Conventional PCR amplification of the nematode-specific D2–D3 region of the 28S rRNA gene was used to verify the presence of “generic” nematode DNA in plant tissue samples. Agarose gel (1.8%) showing amplification products obtained using primers D2A/D3B (Subbotin et al., 2006). Lane 1: 1 kb DNA ladder (GeneRuler 1kb Plus, ThermoFisher Scientific). Lane 2: *Pratylenchus pseudocoffeae* (positive control). Lane 3: nematodes extracted from symptomatic American beech tissue (positive control). Lane 4: mixture of DNA from lanes 2 and 3. Lane 5: naïve American beech *leaf* tissue. Lane 6: naïve leaf tissue spiked with LC. Lane 7: naïve American beech bud tissue. Lane 8: naturally infested *bud* tissue. Lane 9: naïve *bud* tissue spiked with LC. Lane 10: bacterial DNA (negative control). Lane 11: fungal DNA (negative control). Lane 12: PCR blank (molecular-grade water). Lanes 13–15: empty. The presence of double bands likely reflects amplification of multiple nematode taxa within a sample, each contributing different D2–D3 amplicon lengths, as length variation in this region is well documented among nematode species (Subbotin et al., 2006; Pereira and Baldwin, 2016). All observed bands fall within the expected size range (~500–800 bp) and were therefore considered positive for nematode DNA.

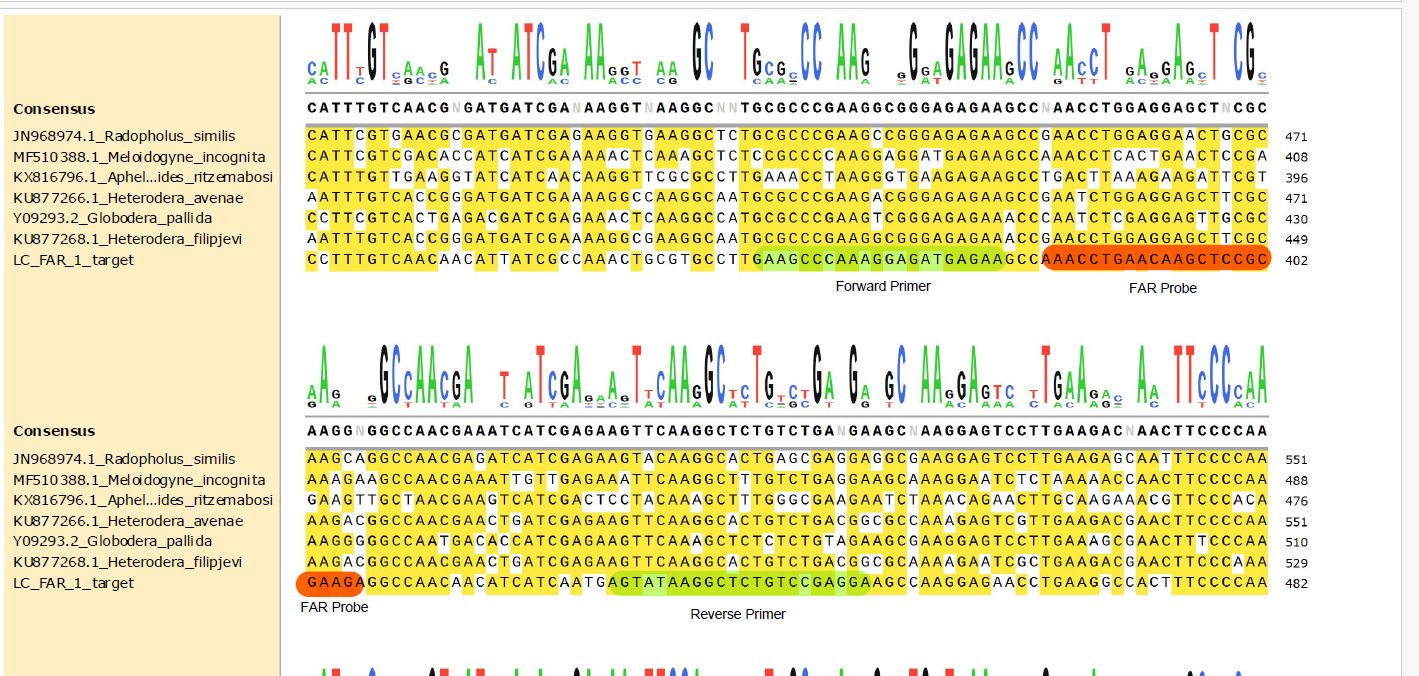

**Figure S3.** Multiple sequence alignment of LC FAR and representative plant-parasitic nematode FAR homologs. Colored boxes indicated the binding sites of the forward primer, hydrolysis probe, and reverse primer.

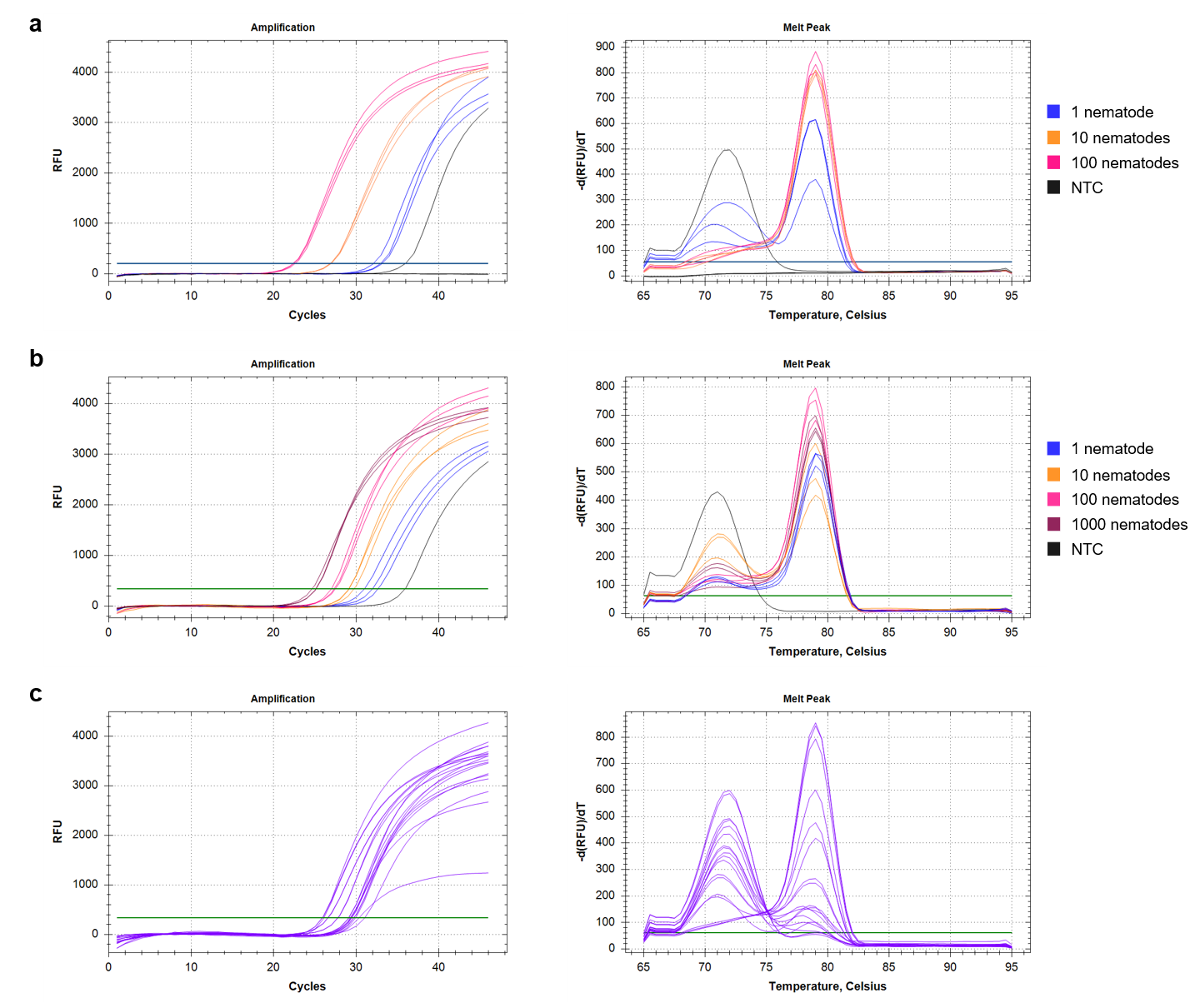

**Figure S4.** SYBR Green qPCR amplification curves and corresponding melt-curve profiles obtained using the FAR primer pair. Panels show results for (a) undiluted DNA extracted from purified nematodes, (b) naïve leaf tissue spiked with LC and diluted 1:15 prior to amplification, and (c) naturally infested leaf tissue samples.

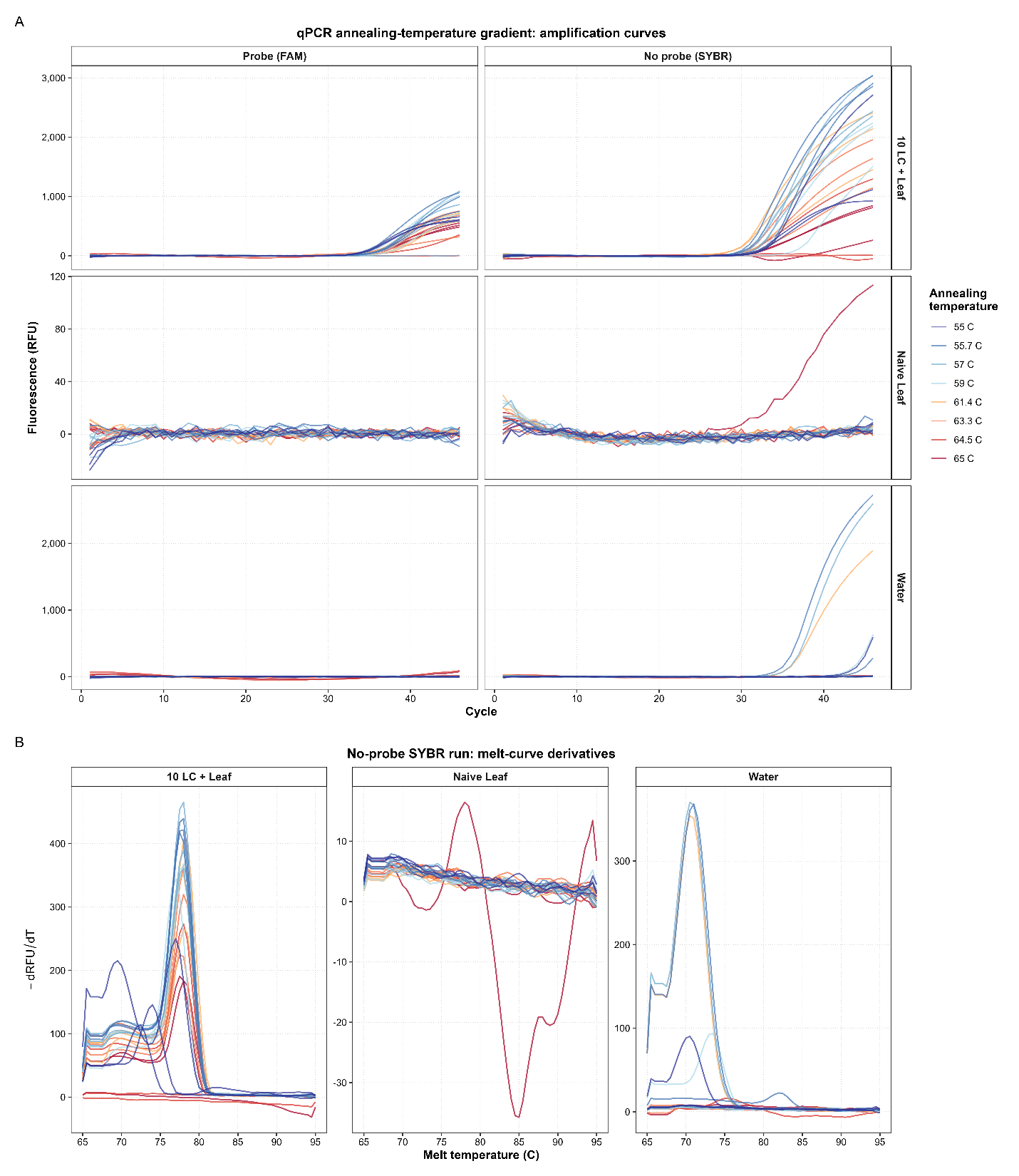

**Figure S5**. Effect of annealing temperature on SYBR Green qPCR amplification and melt curve profile. An annealing temperature gradient (55.0-65.0°C) was evaluated using leaf tissue spiked with 10 LC, naïve American beech leaf DNA, and a non-template water control (water). (A) Amplification curves and (B) corresponding derivative melt curves for each annealing temperature. Curve colors indicate annealing temperature. Although some of the annealing temperatures reduced these nonspecific amplification products, they were not completely eliminated, suggesting the need to develop a hydrolysis probe-based assay to improve assay specificity.

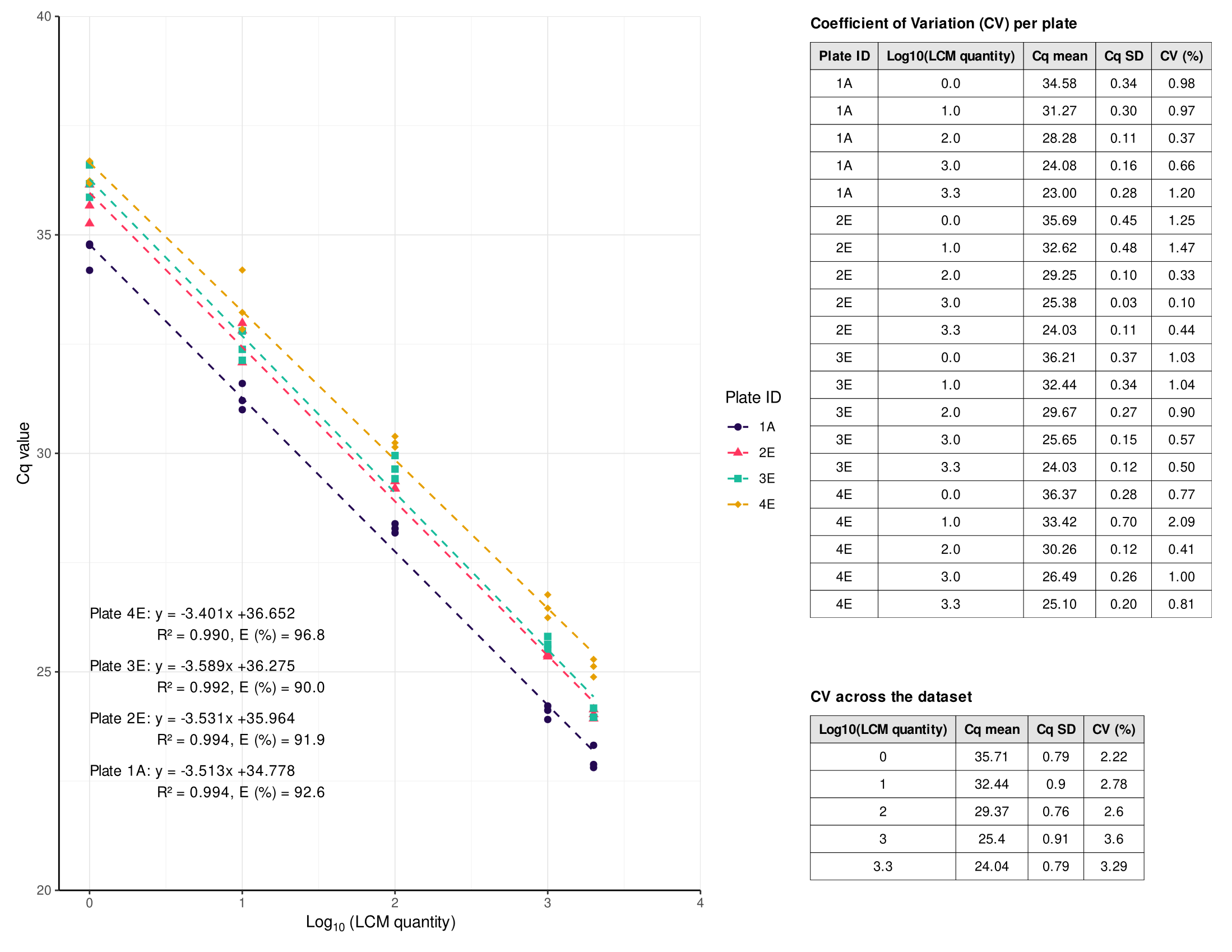

**Figure S6.** Probe-based FAR qPCR standard curves generated from DNA extracted from purified LC are shown for four independent qPCR plates. Each panel displays Cq values plotted against log_10_- transformed nematode quantity, with plate-specific linear regressions, slopes, coefficients of determination (R^2^), and amplification efficiencies. Tables summarize mean Cq values, standard deviations, and coefficients of variation (CV) per concentration and per plate.

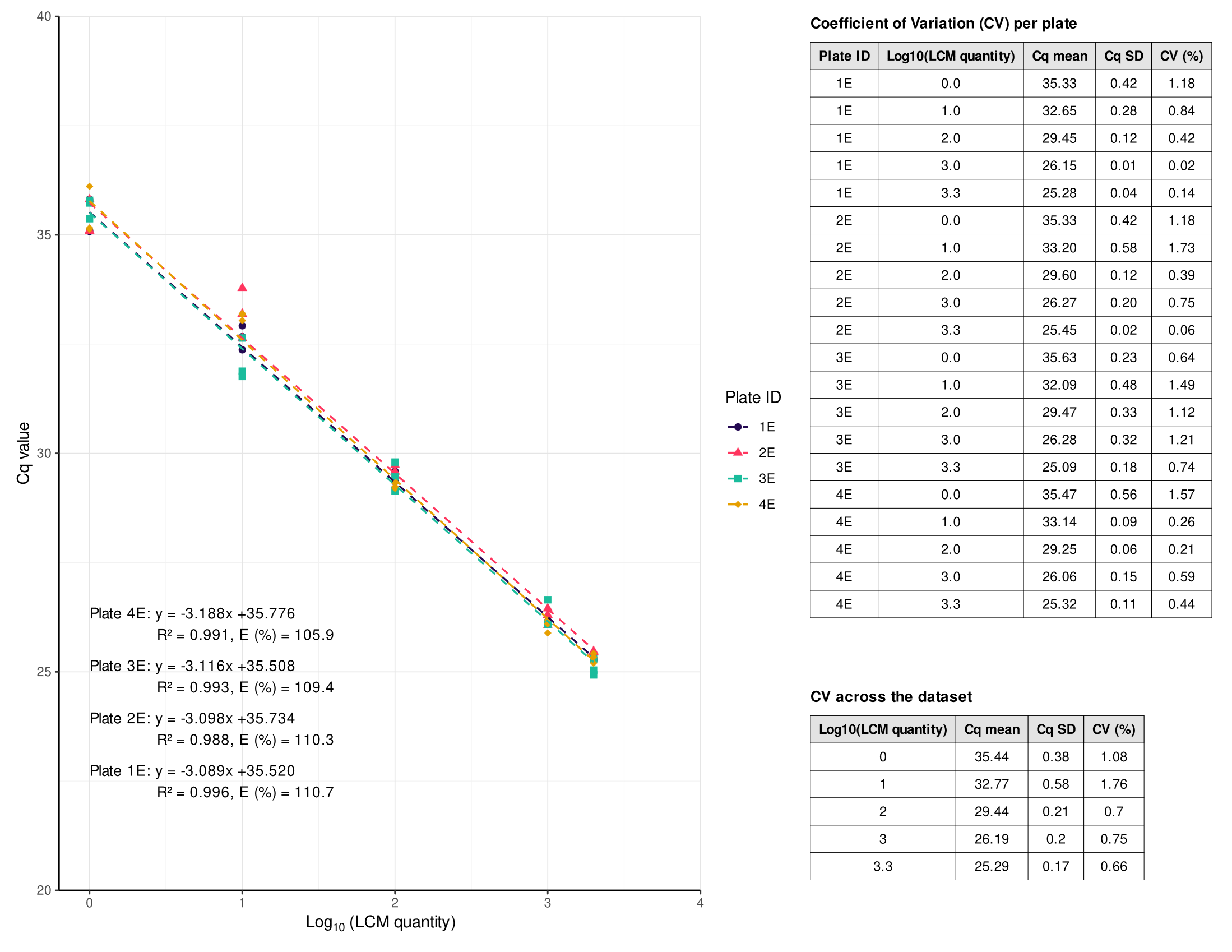

**Figure S7.** Probe-based FAR qPCR standard curves generated from naïve American beech leaf tissue spiked with defined quantities of LC before DNA extraction. Data are shown for four independent qPCR plates. Each panel displays Cq values plotted against log_10_- transformed nematode quantity, with plate-specific linear regressions, slopes, coefficients of determination (R^2^), and amplification efficiencies. Tables summarize mean Cq values, standard deviations, and coefficients of variation (CV) per concentration and per plate.

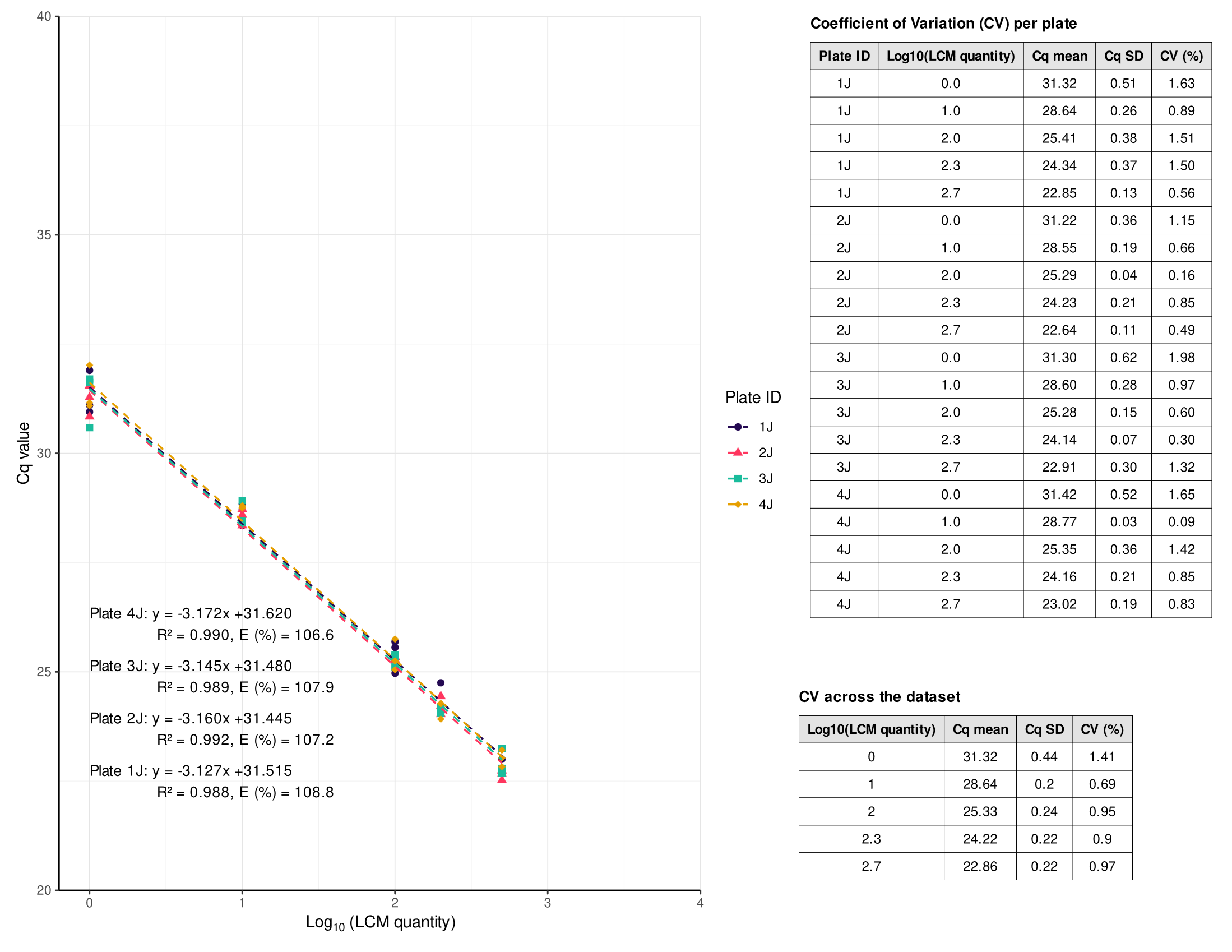

**Figure S8.** Probe-based FAR qPCR standard curves generated from spore-trap tape samples spiked with defined quantities of LC. Data are shown for four independent qPCR plates. Each panel displays Cq values plotted against log_10_- transformed nematode quantity, with plate-specific linear regressions, slopes, coefficients of determination (R^2^), and amplification efficiencies. Tables summarize mean Cq values, standard deviations, and coefficients of variation (CV) per concentration and per plate.

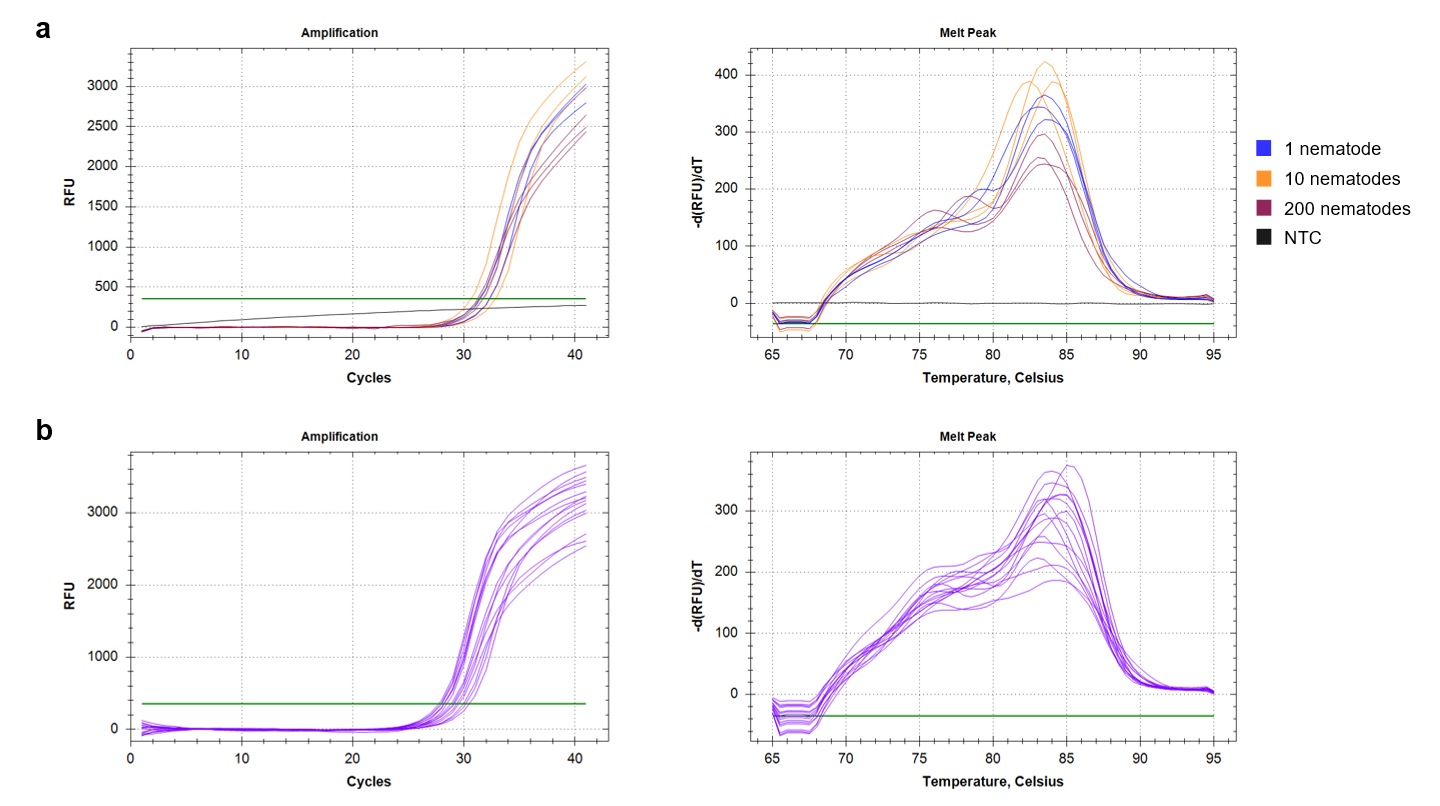

**Figure S9.** Amplification curves (left panels) and corresponding melt-curve profiles (right panels) generated using the ITS-based SYBR Green qPCR assay (Burke et al., 2023) after removal of the extension step from the thermal cycling protocol. (a) Reactions performed with naïve beech leaf tissue spiked with known quantities of LC and (b) naturally infested leaf samples. Melt-curve profiles show multiple peaks and broad peak shapes, indicating non-specific amplification despite protocol modification.

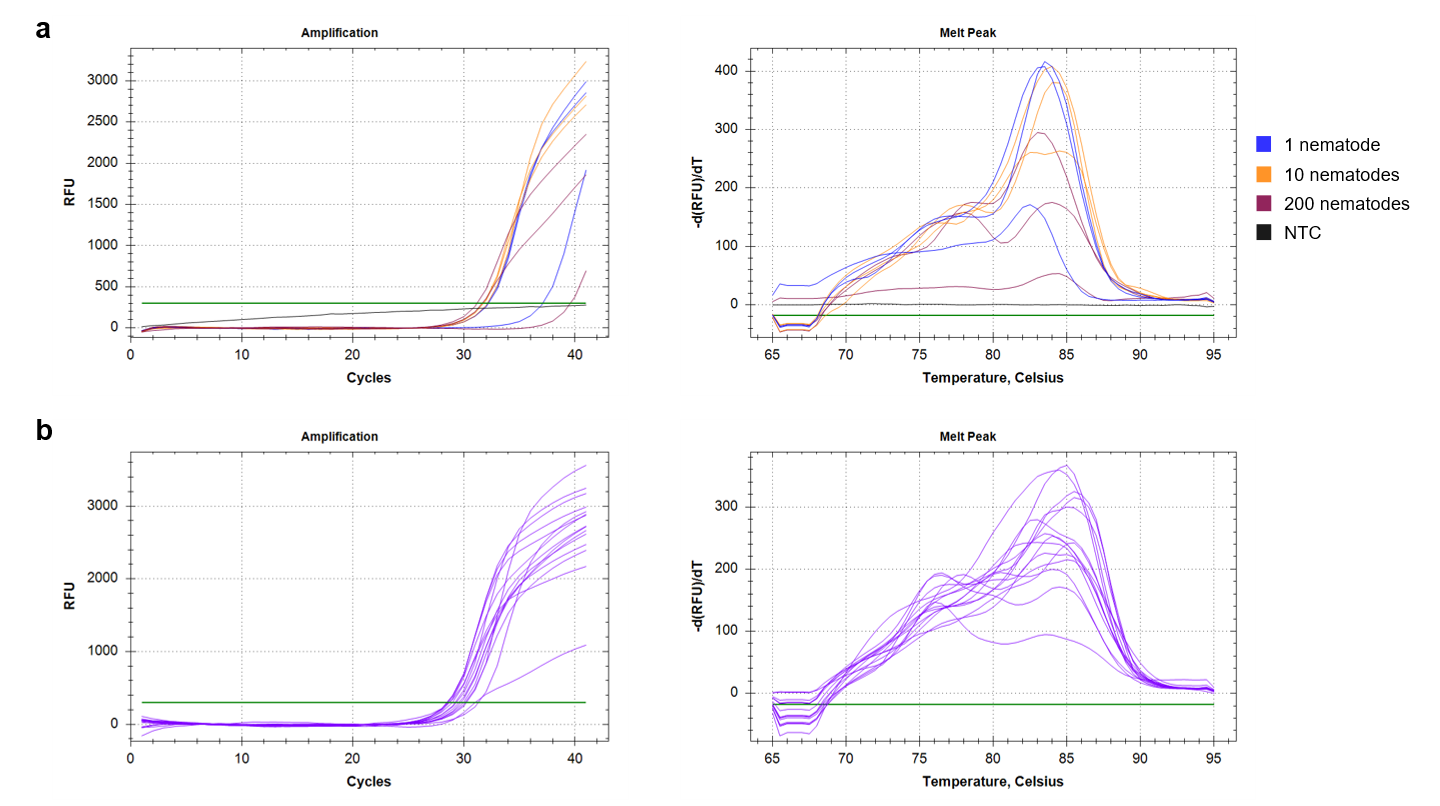

**Figure S10.** Amplification curves (left panels) and corresponding melt-curve profiles (right panels) generated using the ITS-based SYBR Green qPCR assay (Burke et al., 2023) following removal of the extension step and an increase in annealing temperature from 58°C to 59°C. (a) Reactions performed with naïve beech leaf tissue spiked with known quantities of LC and (b) naturally infested leaf samples. Melt-curve profiles show multiple peaks and broad peak shapes, indicating non-specific amplification despite protocol modification.
